# The mitochondrial HSP90 paralog TRAP1 forms an OXPHOS-regulated tetramer and is involved in maintaining mitochondrial metabolic homeostasis

**DOI:** 10.1101/679431

**Authors:** Abhinav Joshi, Joyce Dai, Jungsoon Lee, Nastaran Mohammadi Ghahhari, Gregory Segala, Kristin Beebe, Francis T.F. Tsai, Len Neckers, Didier Picard

**Author notes:** these senior authors contributed equally (Len Neckers, Didier Picard).

## Abstract

**Background:** The molecular chaperone TRAP1, the mitochondrial isoform of cytosolic HSP90, remains poorly understood with respect to its pivotal role in the regulation of mitochondrial metabolism. Most studies have found it to be an inhibitor of mitochondrial oxidative phosphorylation (OXPHOS) and an inducer of the Warburg phenotype of cancer cells. However, others have reported the opposite and there is no consensus on the relevant TRAP1 interactors. This calls for a more comprehensive analysis of the TRAP1 interactome and of how TRAP1 and mitochondrial metabolism mutually affect each other.

**Results:** We show that the disruption of the gene for TRAP1 in a panel of cell lines dysregulates OXPHOS by a metabolic rewiring that induces the anaplerotic utilization of glutamine metabolism to replenish TCA cycle intermediates. Restoration of wild-type levels of OXPHOS requires full-length TRAP1. Whereas the TRAP1 ATPase activity is dispensable for this function, it modulates the interactions of TRAP1 with various mitochondrial proteins. Quantitatively by far the major interactors of TRAP1 are the mitochondrial chaperones mtHSP70 and HSP60. However, we find that the most stable stoichiometric TRAP1 complex is a TRAP1 tetramer, whose levels change in response to both a decline or an increase in OXPHOS.

**Conclusions:** Our work provides a roadmap for further investigations of how TRAP1 and its interactors such as the ATP synthase regulate cellular energy metabolism. Our results highlight that TRAP1 function in metabolism and cancer cannot be understood without a focus on TRAP1 tetramers as potentially the most relevant functional entity.

## Background

Cells adapt their core metabolism in order to sustain survival in an environment where availability of oxygen and nutrients can be limiting [1, 2]. In the past few years, TRAP1, the mitochondrial isoform of the heat-shock protein 90 (HSP90), has been recognized as an important modulator of mitochondrial bioenergetics of normal and cancer cells [3-6]. TRAP1 is directed to the mitochondrial matrix [3, 7, 8] by an N-terminal mitochondrial targeting sequence that is cleaved off upon import [9]. The processed TRAP1 protein is composed of an N-terminal ATPase domain, a middle domain, and a C-terminal dimerization domain; this domain structure is similar to that of cytosolic HSP90 [10], which is the core component of a molecular chaperone machine that is crucial for assisting a large number of “clients” implicated in a wide array of biological processes [11-13]. While cytosolic HSP90 has been extensively studied in the past few decades [13], less is known about the biochemical activities of TRAP1 and how they relate to its role in metabolic regulation (see below). Its crystal structure was recently determined, which has helped to understand its ATPase driven conformational cycle [10, 14-16]. However, in contrast to HSP90, whose ATPase cycle and biological activities are modulated by a large cohort of co-chaperones [13, 17], no co-chaperones have been identified for TRAP1. This may be related to its kinship with bacterial Hsp90, which also functions in the absence of co-chaperones.

TRAP1 expression was found in several studies to be inversely correlated to oxidative phosphorylation (OXPHOS) and OXPHOS-coupled ATP synthesis in different cell types [3, 4]. These data suggested that TRAP1 is a negative regulator of mitochondrial OXPHOS, but the underlying molecular mechanisms have remained controversial. While TRAP1 had been shown to inhibit complex II [4] and IV [3] of the electron transport chain by some, it has also been shown to activate complex II [18] and to support OXPHOS [19] by others. Thus, although TRAP1 has been proposed to play a key role in the induction of the Warburg phenotype of cancer cells, conflicting studies [18, 19] clearly call for additional research to understand how TRAP1 regulates mitochondrial metabolism. A better understanding requires a comprehensive analysis of its interactions with other mitochondrial proteins, in general, and with OXPHOS-associated proteins in particular. Moreover, only a more detailed examination of how TRAP1 and cellular metabolism affect each other will provide sufficient biological insights to evaluate TRAP1 as a potential drug target for the treatment of cancer and other diseases with a metabolic imbalance.

## Results

### Loss of TRAP1 increases OXPHOS due to an anaplerotic increase in glutamine uptake and metabolism

The gene *TRAP1* was disrupted in HEK293T, HCT116, A549, and UMUC3 cells using the CRISPR/Cas9 technology and the workflow presented in Additional file 1: Figure S1a. To confirm that the *TRAP1* knock-out (KO) resulted in an increase in mitochondrial respiration, the cellular oxygen consumption rate (OCR), which is a measure of mitochondrial respiration, was measured in real time in WT and KO HEK293T and HCT116 cells (Fig. 1a, Additional file 1: Figure S1b). Similar to what we had found with mouse adult fibroblasts (MAFs) [3], the KO increases mitochondrial OCR (Fig. 1b) and OXPHOS-linked ATP production (Fig. 1c) in HEK293T cells. An analysis of the energy profile of these cells further showed that although the glycolytic potential of KO cells remained similar to the WT cells (baseline and stressed), the KO made these cells more “aerobic” and dependent upon OXPHOS under normoxic conditions when compared to the WT cells (Fig. 1d). Note that while both HEK293T and HCT116 KO cell lines exhibited increased OCR (Fig. 1a, Additional file 1: Figure S1b), the impact of the KO on OCR is not comparable across the two cell lines, probably because of their different metabolic preferences [20]. The increase in mitochondrial respiration could be suppressed in both HEK293T and HCT116 KO cells by re-introducing TRAP1, but not by overexpressing EGFP directed to the mitochondrial matrix with a TRAP1 mitochondrial targeting signal (MTS) (Fig. 1e, f). The mitochondrial EGFP construct (mitoEGFP) primarily served as a control to verify that overexpression of an unrelated protein in mitochondria did not affect OXPHOS function. Also note that there is always a slight, but statistically insignificant dip in mitochondrial respiration due to transient transfection toxicity (Fig. 1e, f).

**Figure 1.**
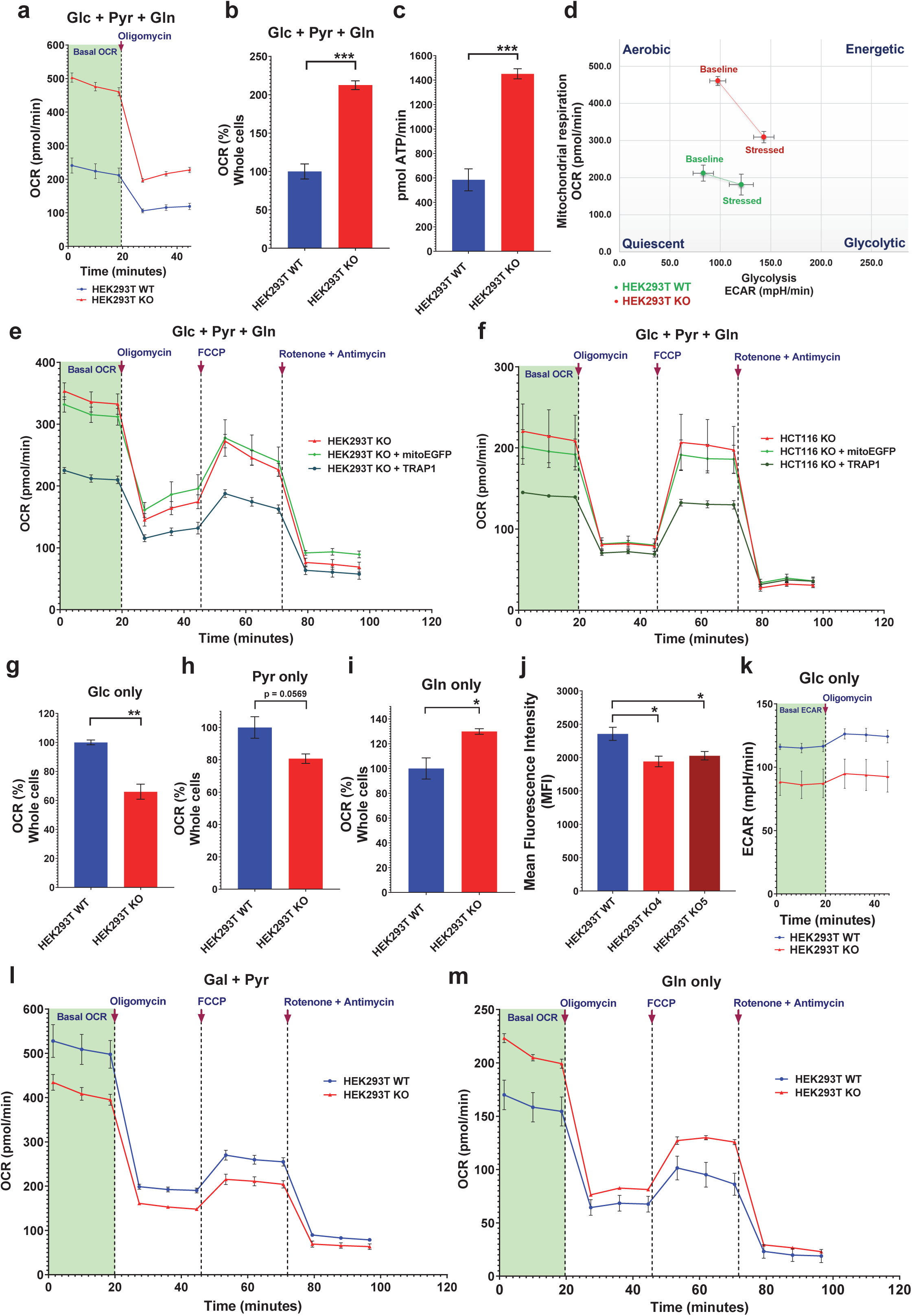
Real-time metabolic profiling of human TRAP1 KO cells. (a) Representative real-time traces of basal OCR of WT and TRAP1 KO HEK293T cells with Glc + Pyr + Gln as carbon sources followed by injection of the ATP-synthase inhibitor (oligomycin at 5 µM) to block mitochondrial respiration. (b and c) Quantitation of basal respiration rates (b) and ATP production (c). ATP production is calculated as (last measurement before oligomycin injection) - (minimum rate measured after oligomycin injection). (d) Comparative energy profiles. The baseline phenotype indicates OCR and ECAR of cells with starting non-limiting assay conditions; the stressed phenotype indicates OCR and ECAR upon exposure to metabolic inhibitors. (e, f) OCR traces with and without the overexpression of TRAP1 or mitoEGFP in HEK293T KO (e) and HCT116 TRAP1 KO (f) cells. The mitochondrial stress test profile is obtained by sequential injection of oligomycin (5 µM), the uncoupler FCCP (2 µM) and the complex I and III inhibitors rotenone (1 µM) and antimycin (1 µM), respectively. (g-i) Comparison of basal OCR of WT and KO HEK293T cells with Glc (g), Pyr (h), and Gln (i) as the only carbon sources. (j) Flow cytometric quantitation of glucose uptake using 2-NBDG (150 µg/ml) with WT and two independent TRAP1 KO HEK293T clones. (k) ECAR traces showing basal glycolytic rates of WT and KO HEK293T cells with Glc as the only carbon source before and after the addition of oligomycin. (l, m) OCR traces of WT and KO HEK293T cells grown in media with Gal + Pyr (l) and Gln (m) as the only carbon sources. All data are reported as means ± SEM (n = 3) with asterisks in the bar graphs indicating statistically significant differences (*p<0.05, **p<0.01, and ***p<0.001).

We next wanted to identify the differential use of carbon sources underlying this respiratory dysregulation. In central carbon metabolism, mitochondrial respiration is primarily driven by the three major carbon sources glucose (Glc), pyruvate (Pyr) and glutamine (Gln). The OCRs of WT and KO cells incubated separately with each of the three carbon sources were determined (Fig. 1g-i).

When grown only on glucose as the primary carbon source, an uptake assay with the fluorescent tracer 2-NBDG showed that HEK293T KO cells have a lower Glc uptake than WT cells (Fig. 1j). As a direct consequence, they display a reduced OCR (Fig. 1g) and rate of extracellular acidification (ECAR), caused by lactate secretion, a measure of the glycolytic flux (Fig. 1k).

To maintain a minimal glycolytic rate and to promote pyruvate oxidation in mitochondria, WT and KO cells were grown overnight in a medium containing galactose and pyruvate (Gal + Pyr) as the only carbon sources [21]. Under these conditions, the ECAR profile tends to mimic the OCR profile (compare Fig. 1l with Additional file 1: Figure S1c and Additional file 1: Figure S1d with Additional file 1: Figure S1e). Real-time respiration monitoring showed that the basal OCR in both HEK293T (Fig. 1l, h) and HCT116 KO cells (Additional file 1: Figure S1d) is decreased, indicating an overall decrease in assimilation of pyruvate into the tricarboxylic acid (TCA) cycle. A separate OCR analysis with only pyruvate as the carbon source gave similar results demonstrating that this outcome was not due to a galactose-induced artefact (Additional file 1: Figure S1f). In contrast, OCR analysis with only Gln as the primary carbon source (Fig. 1m, i and Additional file 1: Figure S1g) indicated a metabolic preference of KO cells for Gln. This may compensate for the reduced Glc or Pyr metabolism and indicate an anaplerotic shift, that is the replenishment of TCA cycle intermediates diverted to various biosynthetic pathways [22], in this case by the increased utilization of Gln. Similarly to Pyr alone, the ECAR profile with only Gln mimicked the OCR profiles of both HEK293T and HCT116 cells, which indicates that Gln is also primarily metabolized in mitochondria in both cell types (Additional file 1: Figure S1h, i).

To confirm the increased Gln uptake and utilization by KO cells, indicated by the OCR experiments, a quantitative flux tracing experiment was performed. For this, isotopically labelled Gln (^13^C-Gln) was added in addition to unlabelled Glc and Pyr as carbon sources (Additional file 2: Figure S2a-c and Additional file 3: Table S1 for absolute quantitation of metabolites; for ^13^C tracing in metabolites, see the NEI area tab in Additional file 4: Table S2). Quantitation of metabolites with increased ^13^C abundance in KO cells are shown in Fig. 2. Both HEK293T and A549 KO cells exhibited a significant increase in total Gln and glutamate concentrations (Fig. 2a), further confirming that KO cells prefer Gln even in the presence of the other two major carbon sources (Glc and Pyr). This is also associated with an increase in the levels of traced TCA cycle intermediates (Fig. 2b) indicating that KO cell metabolism is indeed anaplerotic: the increased Gln uptake and utilization allows the replenishment of TCA cycle metabolites. We further extended this comparison to 42 different quantitated metabolites (Additional file 2: Figure S2 in conjunction with NEI area tab in Additional file 4: Table S2) and also observed a notable increase in ^13^C-traced reduced glutathione (GSH) in both HEK293T and A549 KO cells (Fig. 2c). This may indicate an adjustment to cope with increased reactive oxygen species (ROS), which are often associated with increased OXPHOS [3, 23].

**Figure 2.**
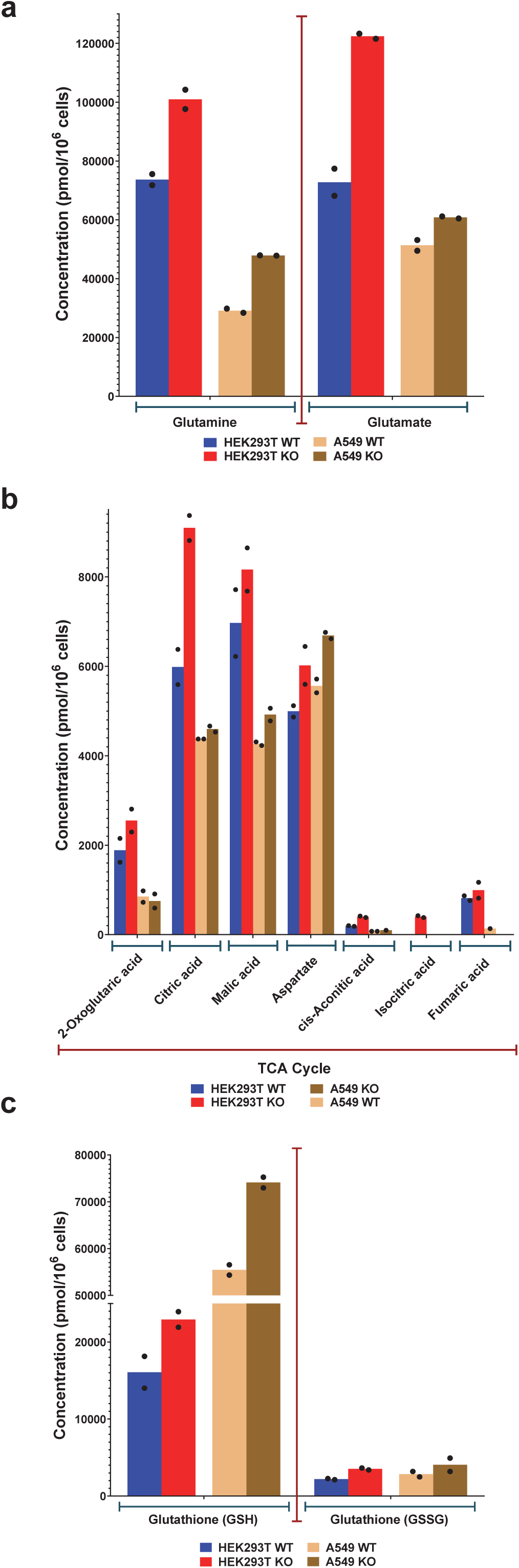
Absolute quantitation of traced metabolites in WT and KO cells. (a-c) Quantitation of total glutamine and glutamate levels (a), TCA cycle metabolites (b), and reduced (GSH) and oxidized glutathione (GSSG) (c) in WT and KO HEK293T and A549 cells. The absolute quantitation shown is for metabolites with increased ^13^C abundance from labelled glutamine (see data in Additional file 4: Table S2). Data points on bar graphs indicate metabolite concentration per 10^6^ cells from each biological replicate (n = 2).

### Full-length TRAP1 but not its ATPase activity is essential to regulate OXPHOS

We next investigated which parts and functions of TRAP1 are necessary to rescue the metabolic phenotype of KO cells. We designed a custom construct to express TRAP1 variants with a C-terminal HA tag and an N-terminal TRAP1-MTS to ensure that proteins are directed into the mitochondrial matrix (Additional file 5: Figure S3a). A mitochondrially targeted EGFP construct (mito-EGFP) was used as a control (Additional file 5: Figure S3b). As mentioned previously, this construct was used to test whether overexpression of an unrelated protein in mitochondria might non-specifically disrupt OXPHOS function (Fig. 1h,i and 3a-d). All TRAP1 truncation mutants as well as the full-length protein were expressed with some exhibiting bands corresponding to precursor proteins with uncleaved MTS and to shorter ones due to N-terminal cleavage (Additional file 5: Figure S3c). The TRAP1 truncation mutants were then overexpressed in the HEK293T KO cells to determine OCR profiles in the presence of all three carbon sources (Fig. 3 a, c). Once again, the OCR data with the mitoEGFP controls confirm a slight reduction in mitochondrial respiration due to transient transfection toxicity (Figs. 1h, i, and 3a, c). However, the slightly lower OCR of cells transfected with the control plasmid expressing mitoEGFP was still significantly higher when compared to the OCR of cells transfected with the WT TRAP1 expression plasmid (Fig. 3 b, d). None of the TRAP1 truncation mutants were able to suppress the KO OXPHOS phenotype to WT levels (Fig. 3 b, d). This indicates that a full-length TRAP1 protein is essential for normal OXPHOS regulation.

**Figure 3.**
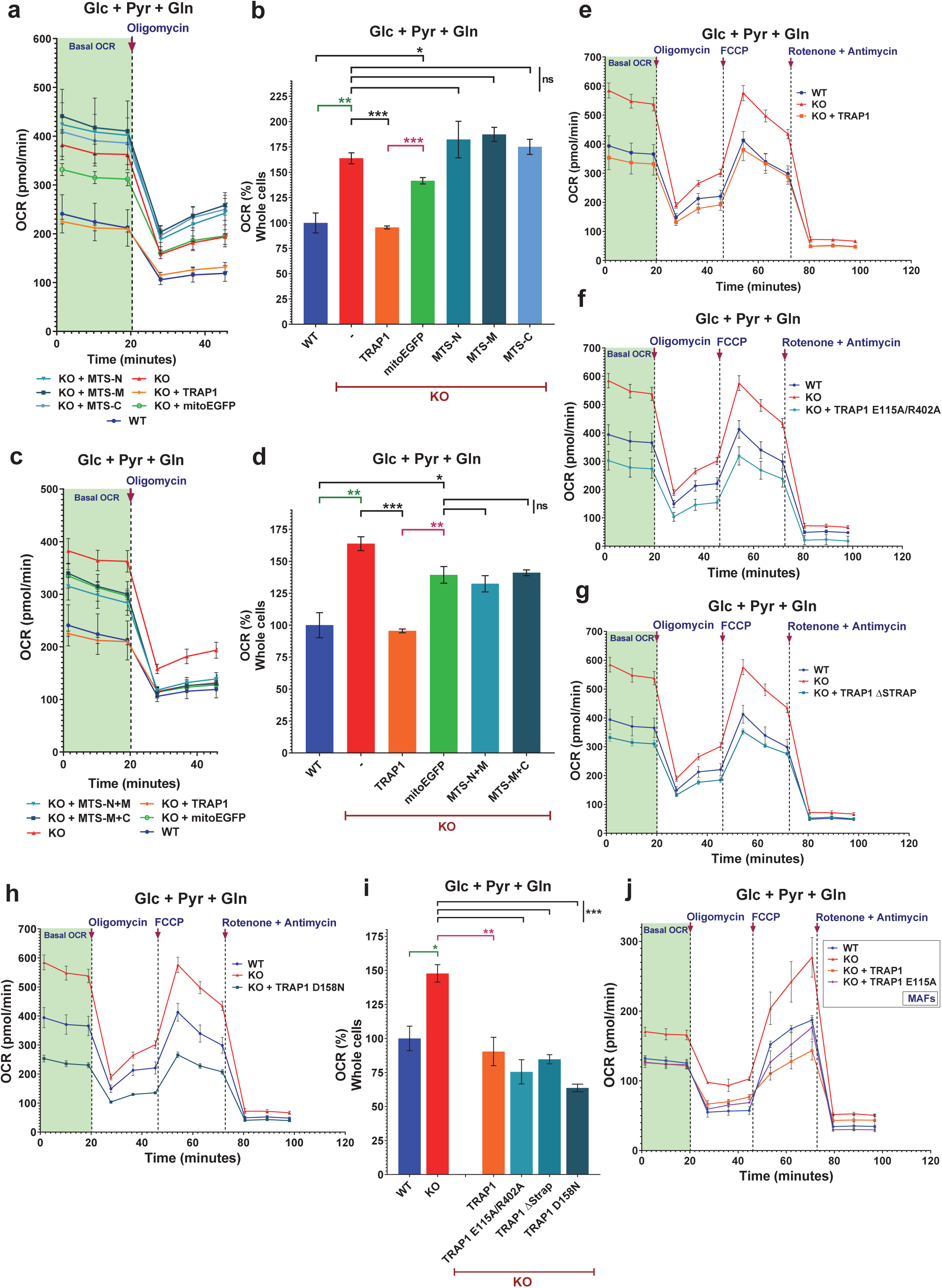
Analysis of the TRAP1 structure activity relationship for metabolic regulation. (a) OCR traces of WT versus KO HEK293T cells exogenously expressing the control proteins mitoEGFP or WT TRAP1, or the TRAP1 truncation mutants MTS-N, MTS-M and MTS-C. (b) Quantitation of the basal respiration rates of WT versus KO HEK293T cells expressing the indicated proteins. (c) OCR traces; experiments as in panel a, but with the TRAP1 truncation mutants MTS-N+M and MTS-M+C. (d) Quantitation of the basal respiration rates of WT versus KO cells expressing the indicated proteins. (e-h) OCR traces of WT versus KO HEK293T cells overexpressing WT TRAP1 (e), the ATPase mutants E115A/R402A (f), ΔSTRAP (g) or D158N (h). (i) Quantitation of the basal respiration rates of WT versus KO HEK293T cells expressing the indicated proteins. (j) OCR traces with WT and KO MAFs and MAF KO cells exogenously expressing either WT TRAP1 or the TRAP1 low ATPase mutant E115A. All data are reported as means ± SEM (n = 3) with asterisks indicating statistically significant differences between compared groups (*p<0.05, **p<0.01, and ***p<0.001).

Since TRAP1 is a paralog of HSP90, a molecular chaperone that is well known to be dependent on its ATPase cycle [24, 25], we speculated that the ATPase activity of TRAP1 might be required for OXPHOS regulation. To test this, we generated a panel of point and truncation mutants that affect this enzymatic activity. Note that our numbering includes the 59 amino acids of the MTS. The following ATPase activity mutants were tested: the double point mutant E115A/R402A with a 10-fold reduced ATPase activity relative to WT (Additional file 5: Figure S3d), the 30-fold hyperactive ATPase mutant ΔStrap, and the moderately activated (2.5-fold) ATPase single point mutant D158N [14]. To our surprise, all ATPase mutants are able to suppress the OXPHOS phenotype of the KO cells, reducing the OCR to WT levels (Fig. 3e-i).

Similar results were obtained when the OCR analysis was done with cells in medium with only Gln as the carbon source (Additional file 5: Figure S3e). We further confirmed the ATPase independence of the complementation by performing a separate real-time OCR analysis with murine cells comparing KO MAFs stably expressing either WT or the single point mutant E115A of human TRAP1 (Fig. 3j).

Note that the mutant E115A was designed by analogy to the yeast HSP90 E33A mutant, which has been reported to be able to bind to ATP, but to be defective for ATP hydrolysis [24, 26]; E115A, similarly to the single mutant mentioned above, binds ATP, but is defective for ATP hydrolysis [15]. Thus, the ability to hydrolyze ATP, at least as well as WT TRAP1, is not essential for the regulation of OXPHOS by TRAP1.

### TRAP1 primarily interacts with other mitochondrial chaperones and OXPHOS-associated proteins

While HSP90 has an exhaustive list of clients and co-chaperones [13, 27-30], the interactome of its mitochondrial paralog remains poorly characterized [6]. After ascertaining that a full-length TRAP1 is essential for OXPHOS regulation, we wondered which proteins interact with TRAP1 and whether these might explain its role in OXPHOS regulation.

We carried out an immunoprecipitation mass spectrometry (IP-MS) experiment with WT TRAP1 and the ATPase mutants E115A/R402A and ΔStrap overexpressed in HEK293T cells (Additional file 6: Figure S4a; Additional file 7: Table S3). To refine this list of identified proteins, the protein interactors were first filtered for validated mitochondrial proteins and then by limiting the dataset to proteins with four or more identified unique peptides. This yielded a list of 82 proteins common to WT TRAP1 and the two ATPase mutants; we took these to represent the most probable TRAP1 interactors (Additional file 8: Table S4). This list primarily contains other mitochondrial chaperones (for example GRP75, CH60, and PHB, which are also known as mtHSP70/mortalin, HSP60, and prohibitin, respectively), OXPHOS complex subunits (ATP synthase, complex I, IV), channel/carrier proteins (TOM/TIM complexes, VDACs) and other mitochondrial enzymes (YMEL1, FAS, ECHA). It is noteworthy that, while we could detect the previously reported TRAP1 interactors SDHA [4, 31], COX4, ATPB, and NDUA9 [19], we did not see others including cyclophilin D [32], PINK1 [33], c-Src [3], HTRA2 [34], and SIRT3 [19] (Additional file 7: Table S3). This may be due to differences in cell lines, relative affinities, interactor-directed IPs, or to other experimental details. More unexpectedly, we did not find any enzymes directly involved in Gln metabolism, such as glutaminase,

glutamine synthase and glutamate dehydrogenase. Note that as a consequence of a decline in Glc and Pyr metabolism, the fluctuating ADP to ATP ratios in KO cells may act as a potent activator of glutaminase to fuel the TCA cycle [35, 36]. ADP has been reported to be the strongest nucleotide activator of glutaminase [35], but ATP, both at low and high concentrations, also stimulates glutaminase activity [36].

For further analysis, we used the total peptide spectral matches (PSM, a metric based on the total number of identified peptides for a given protein), to standardize and to compare the data from IPs with WT and mutant TRAP1. Once standardized to WT, interactors of individual TRAP1 mutants could be compared amongst themselves, and as a ratio to the respective TRAP1 versions (set to 100). It is striking that TRAP1 interacting proteins segregate into two major groups based on how much protein was pulled down with WT or mutant TRAP1 (Fig. 4a, Additional file 8: Table S4). Quantitatively, the mitochondrial chaperones GRP75 (mtHSP70), CH60 (HSP60) and PHB2 are the main TRAP1 interactors while all other interactors segregate into the second less abundant group (Fig. 4a, inset).

**Figure 4.**
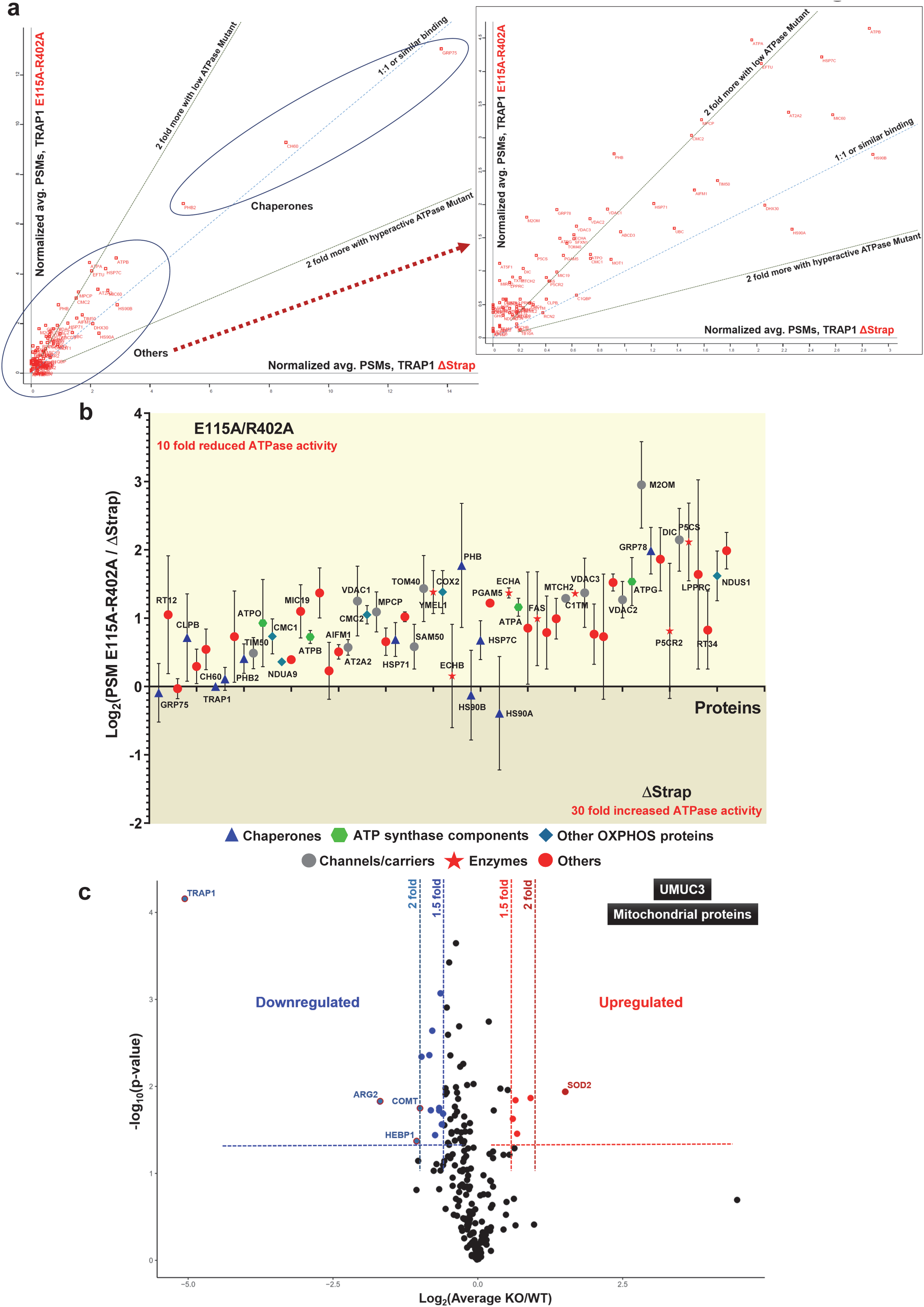
TRAP1 proteomics. (a) Comparative relative abundance of proteins immunoprecipitated with the indicated TRAP1 ATPase mutants. The scatter plot was constructed with an average of normalized PSM values (TRAP1 itself was set to 100) to compare the interactomes of TRAP1 mutants E115A/R402A with low ATPase activity (Y-axis) and the hyperactive ATPase mutant ΔStrap (X-axis); the bigger the distance from the origin on either axis, the more binding there is. The dashed red arrow connects the area near the origin of the plot to the zoomed out inset. (b) Relative quantitation of protein binding to the TRAP1 mutants E115A/R402A and ΔStrap based on log_2_ ratios of normalized PSM values. Proteins above the X axis interact more with the mutant E115A/R402A than the mutant ΔStrap. (c) Volcano plot showing up- or downregulated mitochondrial proteins in a comparison of WT and TRAP1 KO UMUC3 cells. These data are based on the SILAC analysis of the whole cell proteome filtered for mitochondrial proteins.

Consistent with what has been observed for yeast HSP90 by a two-hybrid screen [37], most of the TRAP1 interactors, except the major mitochondrial chaperones mtHSP70 (GRP75) and HSP60 (CH60), have a preference for binding the TRAP1 mutant E115A/R402A, which has a 10-fold reduced ATPase activity and might therefore accumulate in the ATP-bound conformation (Fig. 4b, Additional file 8: Table S4). This preference for the ATP-bound state could also be seen when low and hyperactive ATPase mutants were individually compared to WT TRAP1 (Additional file 6: Figure S4b, c).

Taken together, these results show that while the ATPase activity of TRAP1 can vary greatly without affecting OXPHOS regulation and interaction with other mitochondrial chaperones, TRAP1 ATPase activity is inversely correlated with binding to other TRAP1 interactors.

### Loss of TRAP1 has a minor impact on mitochondrial and total cellular proteomes

We speculated that the absence of TRAP1 might destabilize some of its direct or indirect interactors or lead to a compensatory upregulation of other proteins. We used two separate approaches to identify such proteome changes. First, we performed a quantitative SILAC MS analysis comparing WT to KO UMUC3 cells. 200 mitochondrial proteins were detected (Additional file 9: Table S5). Among this group of interactors, we found little variation comparing KO to WT cells when the minimum significant fold change is set to 2 (p<0.05) (Fig. 4c). Even with a cutoff of 1.5-fold, only a few alterations in the mitochondrial proteome could be seen (Fig. 4c, Additional file 9: Table S5). With the notable exception of PHB2 (when a 1.5-fold change is set as threshold), most of the mitochondrial proteins including those predicted to interact with TRAP1 (especially the subunits of the ATP-synthase complex highlighted by the analysis of Fig. 4b), show no significant up- or downregulation in UMUC3 KO cells (Additional file 9: Table S5). Thus, the TRAP1 KO does not have a significant impact on the stability of the mitochondrial proteome.

Second, we did a label free quantitation (LFQ) MS analysis of the total cellular proteome with WT and KO HEK293T and HCT116 cells cultured with the three different cocktails of carbon sources (Glc + Pyr + Gln, Gal + Pyr only, Gln only; Additional file 10: Table S6). We reduced the initial list of 4578 proteins to 2660 proteins by using as criterion the identification of at least seven unique peptides per protein (Additional file 11: Table S7). The comparison of the LFQ^KO^/LFQ^WT^ ratios for these proteins from cells cultured in medium with all three carbon sources did not reveal any significant changes (Additional file 6: Figure S4d, e). Although a few proteins were observed outside the 2-fold limit, they were not consistent across HEK293T and HCT116 cells to warrant a correlation with the loss of TRAP1. The LFQ ratio profiles turned out to be similar for media with other combinations of carbon sources (Additional file 11: Table S7).

*In toto*, all three MS experiments indicated that while TRAP1 interacts with multiple mitochondrial proteins, its loss does not have much of an impact on the mitochondrial or cellular proteomes.

### TRAP1 forms an oligomeric complex

Our IP-MS experiment suggested that TRAP1 associates with a number of proteins of the mitochondrial matrix in a manner independent of its own ATPase activity. To explore this further, we decided to separate mitochondrial extracts made with a non-ionic detergent from HEK293T cells on clear native polyacrylamide gels (native PAGE) capable of resolving molecular complexes between 1 MDa and and 240 kDa (Fig. 5). We chose clear native PAGE rather than blue native gels because the milder conditions can better preserve the structural and functional integrity of protein complexes; overall, despite the slightly poorer resolution compared to blue native gels, clear native gels have been demonstrated to yield largely comparable results, notably for mitochondrial complexes [38]. We expected the migration of complexes with a protein such as TRAP1 with a pI of 6.40 in a separating gel at pH 8.8 to be reasonably well correlated with molecular weight and size. When blotted for endogenous TRAP1, a single molecular complex of ∼300 kDa could be seen, which is absent from KO cells (Fig. 5). However, the molecular weight of the detected complex was not exactly what was expected if a TRAP1 dimer was in a complex with mtHSP70, HSP60 or even both proteins. Moreover, looking at overexpressed WT or ATPase mutant TRAP1 side by side, we found that the E115A/R402A mutant forms a complex of the same size as WT TRAP1 whereas the hyperactive ATPase mutant (ΔStrap) seems to form a slightly larger or conformationally different, more slowly migrating complex (Fig. 5).

**Figure 5.**
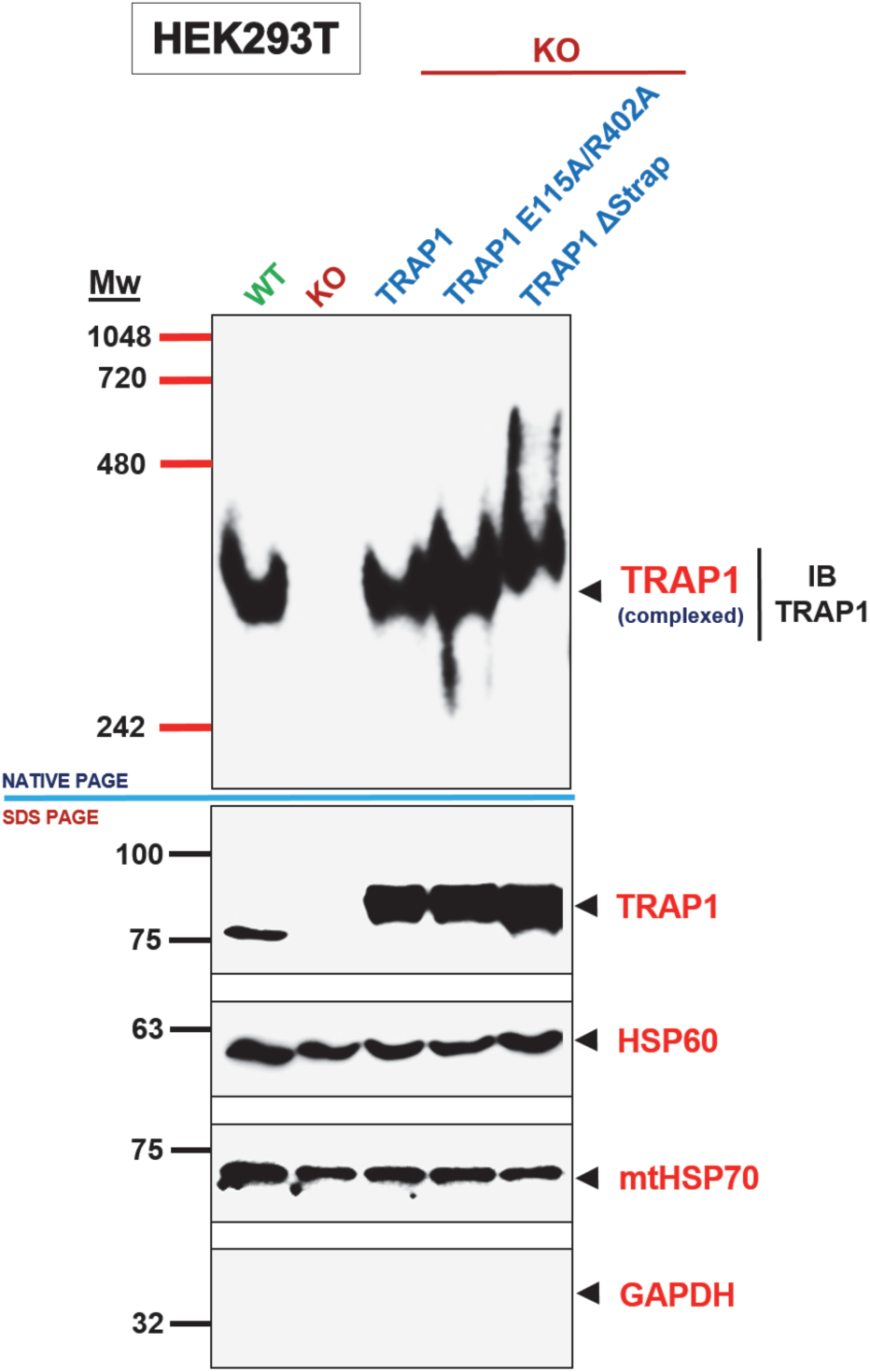
TRAP1 exists as a complex in mitochondria. Immunoblot of a native protein gel (NATIVE PAGE) showing TRAP1 complexes in mitochondrial extracts of WT versus KO HEK293T cells, and KO cells overexpressing WT TRAP1 or the TRAP1 mutants E115A/R402A or ΔStrap. Note that the ΔStrap mutant forms a slightly larger complex when compared to the others. The immunoblot was probed with a TRAP1 antibody. A parallel immunoblot was performed on the same samples under denaturing conditions (10% SDS PAGE) to check the expression levels of TRAP1. HSP60 and mtHSP70 were used as positive and GAPDH as negative controls to check the quality of the mitochondrial extracts.

To determine what the 300 kDa TRAP1 complex contains, we expressed a TRAP1-GST fusion protein and GST alone as a negative control, and applied the workflow described in Additional file 12: Figure S5A for a GST-pulldown MS analysis. Upon setting the cutoff for an interactor at a minimum of eleven unique peptides, no mitochondrial chaperone could be detected in the excised gel piece. Apart from TRAP1, only proteins that were also co-purified with GST alone could be identified (Additional file 12: Figure S5b, Additional file 13: Table S8). Hence, the high molecular weight TRAP1 complex (∼400 kDa in the case of TRAP1-GST) only contains TRAP1-GST. The TRAP1 interactors mtHSP70 and HSP60 may not be sufficiently stably bound to remain associated during native gel electrophoresis. The sizes of the TRAP1 and TRAP1-GST complexes are consistent with TRAP1 forming a stable tetramer or a dimer of dimers.

### The TRAP1 complex is induced in response to OXPHOS perturbations

Based on the hypothesis that an oligomerized TRAP1 complex might be the functional entity of TRAP1, we checked its levels when OXPHOS is inhibited with a prolonged exposure of HEK293T cells to hypoxia in various media (Fig. 6a). Although the baseline levels of the TRAP1 complex vary in cells adapted to different carbon sources in normoxia (left part of Fig. 6a), we saw a consistent increase in the levels of the TRAP1 complex when cells were placed in hypoxia. It is notable that the maximum increase in the levels of the TRAP1 complex was observed with cells grown in Gal + Pyr medium when they were exposed to hypoxia (Fig. 6a). Cells with this carbon source combination exclusively rely on OXPHOS for respiration (Additional file 1: Figure S1, compare panels d and e). Considering that the ATP synthase is one of the major OXPHOS complexes that is inhibited by prolonged hypoxia [39] and that we had found ATP-synthase components to be amongst the main TRAP1 interactors (see Fig. 4b), we asked whether inhibition of the ATP-synthase complex would affect TRAP1 oligomerization (Fig. 6b). To this end, we compared the levels of the TRAP1 complex from HEK293T cells exposed to hypoxia or to the ATP-synthase inhibitor oligomycin under normoxic conditions. Under hypoxic conditions, the induction of the TRAP1 complex is slow and only seems to initiate around 6 hrs. (Fig. 6b). The slow time course may reflect the slow depletion of oxygen from the medium and cells rather than a characteristic of mitochondria or the TRAP1 complex. There is also an overall increase in the levels of TRAP1 protomers in cells exposed to hypoxia (Fig. 6b, middle panel with SDS-PAGE), but this induction does not appear to be HIF1α-mediated (Additional file 14: Figure S6a). In contrast, oligomycin induces a more rapid accumulation of the TRAP1 complex above basal level without a noticeable concomitant increase in total TRAP1 protein levels (Fig. 6b).

**Figure 6.**
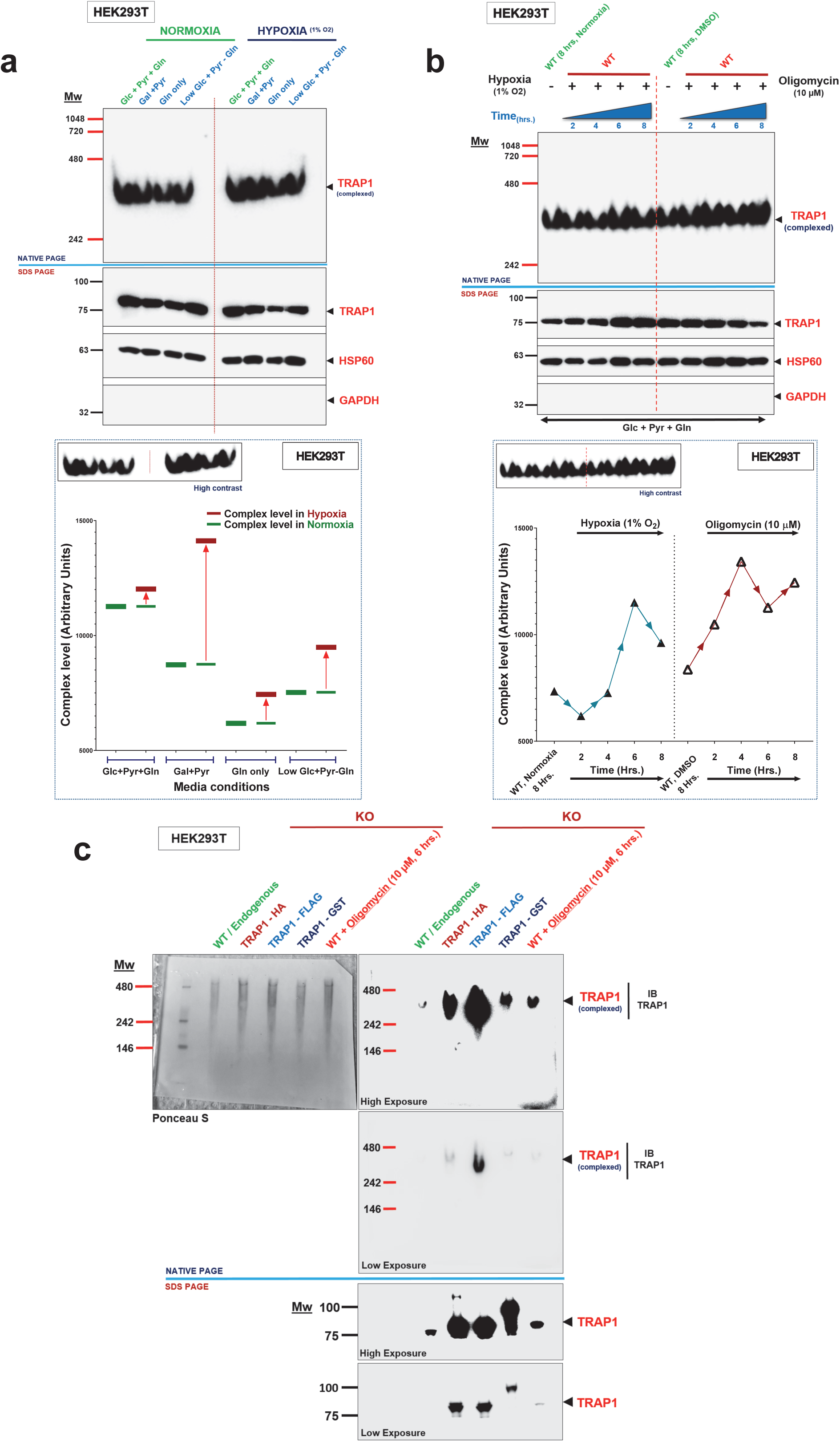
The TRAP1 complex is induced by OXPHOS inhibition. (a) Immunoblot of a native gel analysis of TRAP1 complexes from HEK293T cells grown with different carbon sources under normoxia or hypoxia (1% O_2_) overnight. Lower panel: graphical representation of the levels of the TRAP1 complex shown in the upper panels; band intensities were quantitated using ImageJ. (b) Immunoblot of a native gel analysis of TRAP1 complexes from cells subjected to hypoxia (1% O_2_) or an oligomycin (10 µM) treatment in parallel (in normoxia) for 2, 4, 6 and 8 hrs. The lower panel shows the quantitation. (c) Immunoblot of a native gel analysis to compare the complexes formed by endogenous TRAP1 and the indicated overexpressed tagged versions of TRAP1. For comparison, the endogenous TRAP1 complex was induced with oligomycin (10 µM). Note that no TRAP1 dimer is detectable at steady state under any condition. All native gel immunoblots were probed with a TRAP1 antibody and a parallel immunoblot under denaturing conditions (7.5% SDS PAGE) was also performed to check TRAP1 levels. HSP60 and GAPDH served as positive and negative controls to check the quality of the mitochondrial extracts. All quantitations with ImageJ shown are for a single native gel; similar results were obtained in 3 independent experiments.

Our results showing the existence of a previously unreported TRAP1 oligomeric complex were quite surprising considering that structural [10, 15] and crosslinking [40] studies had only reported TRAP1 to exist as a dimer. To determine whether the dimer and tetramer co-exist at steady state without crosslinking, we compared the endogenous TRAP1 to our panel of full-length TRAP1 proteins with different tags using native gel analysis capable of resolving complexes from 480 kDa to ∼120 kDa (Fig. 6c). Although all protomers were well expressed (Fig. 6c, lower panel with SDS-gels), we did not observe any TRAP1 dimer at steady state, neither with endogenous TRAP1 nor upon further induction of the TRAP1 complex with oligomycin (Fig. 6c). This suggests that a TRAP1 tetramer and not a dimer is the functional unit of TRAP1 in mitochondria.

All of the experiments presented so far regarding the TRAP1 complex were performed solely with HEK293T cells. We therefore confirmed the existence and inducibility of the TRAP1 complex in four other cell lines: breast cancer-derived cell lines MCF-7 and MDA-MB-134, the prostate cancer cell line PC3, and the colon cancer cell line HCT116. A high molecular weight TRAP1 complex, which is rapidly induced further in response to ATP synthase inhibition, was readily detected in each cell line (Additional file 14: Figure S6b).

Next, we assessed the impact of inhibitors of the electron transport chain (ETC) on the TRAP1 complex in MCF-7 and HEK293T cells (Fig. 7a and Additional file 15: Figure S7). Both cell lines showed an accumulation of the TRAP1 complex when the ATP synthase was compromised (Fig. 7a and Additional file 15: Figure S7). In contrast to the inhibition of the ATP-synthase complex (complex V of the ETC), inhibition of complexes I or III or both reduced the TRAP1 complex levels in both cell lines (Fig. 7a and Additional file 15: Figure S7). Therefore, we tested whether inhibition of ATP synthase could override the effects of complex I and III inactivation (Fig. 7b). This was examined at the 3 and 6 hr time points with a combination of rotenone + antimycin and oligomycin + rotenone + antimycin in parallel. Indeed, inhibition of ATP synthase was able to override the suppressive effect of the combined inhibition of complexes I and III on the TRAP1 complex in HEK293T cells, as can be most clearly seen at the 6 hr time point (Fig. 7b).

Having found that the levels of the TRAP1 complex change upon inhibiting OXPHOS, we wondered what would happen if OXPHOS were upregulated. This question is not trivial to address experimentally as it appears that most cells in culture operate OXPHOS at or close to maximal capacity. We decided to culture HEK293T cells on glucose as the only carbon source and then to force them to divert pyruvate to OXPHOS by blocking its conversion to lactate with a lactate dehydrogenase inhibitor (LDHi) (Fig. 7c). This treatment increased the basal OCR of HEK293T cells by more than 2-fold compared to the low basal value of cells grown with glucose as the only carbon source (Fig. 7d). When the cells were treated for 2, 4 or 6 hrs with the LDHi under this condition, we observed a steady increase in the induction of the TRAP1 complex (Fig. 7e). Thus, the TRAP1 complex can be induced both in response to inhibition of OXPHOS at the level of ATP synthase and to an increase of OXPHOS.

**Figure 7.**
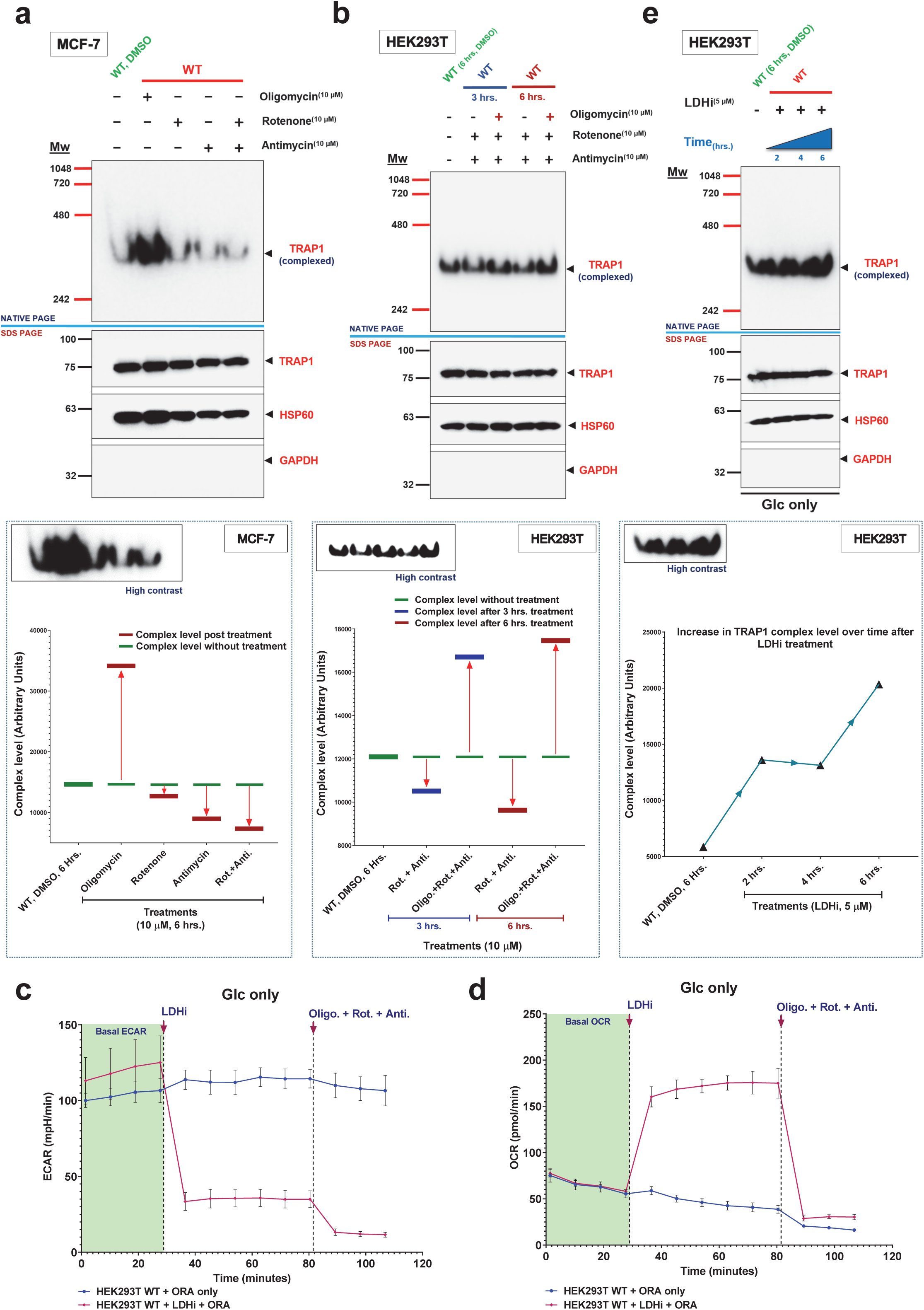
TRAP1 complexes with inhibition and induction of OXPHOS. (a) Immunoblot of a native gel analysis of TRAP1 complexes from MCF-7 cells upon inhibition of OXPHOS at different steps. The lower panel shows the quantitation. (b) Immunoblot of a native gel analysis of TRAP1 complexes from HEK293T cells upon inhibition of OXPHOS at different steps alone and in combination. (c) ECAR profiles of HEK293T cells treated with OXPHOS inhibitors (ORA, cocktail of oligomycin, rotenone and antimycin) with or without an LDH inhibitor (LDHi, 5 µM). (d)OCR profile of HEK293T cells treated with OXPHOS inhibitors (ORA, cocktail of oligomycin, rotenone and antimycin) with or without an LDH inhibitor (LDHi, 5 µM). (e)Immunoblot of a native gel analysis of TRAP1 complexes from HEK293T cells treated with LDHi for 2, 4 and 6 hrs.

## Discussion

The role of TRAP1 in the regulation of mitochondrial metabolism had remained controversial. Here we provide new insights that should help clarify the impact of TRAP1 on cellular energy metabolism and, conversely, on how changes in cellular metabolism affect TRAP1 itself. In most cell lines grown in rich medium, the primary phenotype of a loss of TRAP1 function is an increase in mitochondrial respiration and ATP production [6]. Based on a limited metabolomics analysis we had previously speculated that the increase in OXPHOS in TRAP1-deficient cells is anaplerotic [3]. By using CRISPR/Cas9-generated *TRAP1* KO cell lines, OCR experiments with restricted carbon sources, and metabolomics, we have discovered that the increase in OXPHOS in TRAP1 KO cells is a consequence of stimulated Gln metabolism. The anaplerotic metabolism of TRAP1 KO cells itself might be a compensatory response to a decline in glucose uptake and pyruvate assimilation into the TCA cycle under normoxic conditions. Why cellular glucose uptake and mitochondrial pyruvate utilization are reduced in the absence of TRAP1 remains to be elucidated. Interestingly, the increase in Gln metabolism of TRAP1 KO cells is also channeled into the synthesis of GSH, possibly to buffer the increased ROS produced as a consequence of upregulated OXPHOS [3, 31, 41].

Surprisingly, we could not find any evidence of an interaction between TRAP1 and the enzymes directly involved in Gln metabolism in our TRAP1 IP-MS data, even though we had observed that TRAP1 KO cells grown in Gln only medium are more sensitive to the glutaminase inhibitor CB-839 than WT cells (data not shown).Glutaminase activity has previously been shown to be stimulated by ATP [36], and even more strongly by ADP [35]. Therefore, we speculate that the increase in the ADP/ATP ratio associated with the decline in glucose and pyruvate metabolism in TRAP1 KO cells provides a strong stimulus for activation of mitochondrial glutaminase resulting in a re-equilibrated ADP/ATP ratio. Unfortunately, there is at present no experimental means to measure glutaminase activity in real time as a function of ADP or ATP levels in live cells.

Our efforts to understand how TRAP1 functions as a negative regulator of mitochondrial OXPHOS in normoxia show that the restoration of properly regulated, wild-type levels of OXPHOS requires full-length TRAP1. While this is not surprising, it was unexpected that the ATPase activity of TRAP1 does not correlate with its ability to restore OXPHOS to WT levels. This finding strongly suggests that the ATPase activity of TRAP1 is not essential for OXPHOS regulation. This is reminiscent of relatively recent findings with cytosolic HSP90 indicating that the rate of ATP hydrolysis does not correlate with the ability of this molecular chaperone to support yeast viability [42], while ATP binding is absolutely essential [24, 26, 42].

Similarly, some activities of the bacterial form of HSP90, HtpG, do not depend on its ATPase activity [43]. In the case of TRAP1, it was not possible to test whether ATP binding *per se*, even without hydrolysis, is essential for TRAP1 to regulate OXPHOS. As of today, there is no TRAP1 point mutant that is functionally equivalent to the yeast HSP90 mutant D79N, which abolishes ATP binding [24, 26]. Studies on substitutions of D158, the corresponding amino acid of TRAP1, have yielded conflicting results [14, 44], although the observation that the ATPase activity of D158N is several fold greater than that of WT [14] implicitly proves that this particular mutant can still bind ATP.

Whereas the rate of TRAP1 ATP hydrolysis does not influence its role in OXPHOS regulation, the TRAP1 IP-MS data described in this study show that the ATP hydrolysis rate does affect TRAP1 interactions with other non-chaperone proteins. While the binding of major TRAP1 interactors such as the molecular chaperones mtHSP70 and HSP60 remains unaffected by the ATPase activity of TRAP1, the binding of most non-chaperone interactors, similarly to what has been reported for cytosolic HSP90 interactors [42], is inversely correlated with TRAP1 ATPase activity.

Cytosolic HSP90, with its large clientele of proteins, is a major network hub in the cellular proteome; as a result, pharmacological inhibition of HSP90 greatly destabilizes the cellular proteome [45-50]. This is in stark contrast to what we found for TRAP1, whose loss does not cause a significant imbalance in either the mitochondrial or cellular proteomes. Even the highest confidence TRAP1 interactors such as ATP synthase remain unaffected. Probably the most notable change in TRAP1 KO cells is the increase in mitochondrial SOD2 protein levels. This may be a secondary response to the increase in GSH levels to reduce the oxidative stress that is a direct consequence of increased OXPHOS in TRAP1 KO cells.

Since the major goal of this study was to understand how TRAP1 regulates OXPHOS, we chose to focus on TRAP1 interactors that did not differentially segregate between the ATPase mutants in our IP-MS analysis. This category of interactors includes mtHSP70 and HSP60 as the main interactors of TRAP1. Since cytosolic and bacterial HSP90 work as a chaperone machine in the cytosol with the HSP70/HSP40 system [51, 52], we set out to investigate and to visualize such complexes for TRAP1 by native PAGE. The TRAP1 complex that we saw had an unexpected apparent molecular weight close to 300 kDa. If TRAP1 were to associate with HSP60 alone, this complex should have been ≥ 570 kDa in size since TRAP1 has been reported to form a dimer [10, 15, 53], and since the minimum functional unit of HSP60 is reported to be an oligomerized heptamer [54]. As a heterotetramer with mtHsp70, it could have been close to the observed size of 300 kDa [16]. However, the MS analysis of proteins pulled down with a TRAP1-GST fusion protein revealed that the detected TRAP1 complex is composed solely of TRAP1. Considering the apparent size of the 400 kDa TRAP1-GST complex, we concluded that it must be composed of four TRAP1 protomers, organized either as a tetramer or as a dimer of dimers. Previous biochemical and structural analyses with purified recombinant TRAP1 had shown that TRAP1 exists as a homodimer [10, 40]. Despite our specific efforts to detect them by native PAGE, both for endogenous and overexpressed TRAP1, we were unable to do so. Thus, TRAP1 might not functionally exist as a stable dimer in the mitochondrial matrix at steady state, but only as the proposed tetramer. Intriguingly, higher order structures for cytosolic HSP90 have been found upon exposure to elevated temperatures [55-57]. Moreover, bacterial HtpG was found to be composed of dimers of dimers in the crystal structure [58]. While it remains unclear whether these structures are physiologically relevant for either eukaryotic or bacterial HSP90, our results indicate that they may well be for TRAP1 in mitochondria, which have been demonstrated to function at a higher temperature than the cytosol [59]. Future biochemical and structural analyses of TRAP1 complexes isolated from mitochondria or formed *in vitro* could help to characterize the determinants of this higher order assembly.

In view of the evidence that a TRAP1 tetramer may be the primary “functional unit” of TRAP1, we reasoned that its levels might be influenced by fluctuating OXPHOS. Indeed, when we inhibited OXPHOS by exposure of cells to hypoxia, we observed that the levels of the TRAP1 complex increased with a corresponding increase in the total mitochondrial protomer levels as observed with native and denaturing PAGE, respectively. However, this increase in TRAP1 complex and total protomer levels cannot be attributed to HIF1α as its overexpression does not induce TRAP1 mRNA expression. Hypoxia is a strong inhibitor of ATP synthase [39, 60] and thus, the induction of the TRAP1 complex can be observed both upon inhibiting ATP synthase by hypoxia and in normoxic cells with the pharmacological inhibitor oligomycin. The connection with the ATP synthase is further supported by our finding that multiple subunits comprising the ATP-synthase complex interact with TRAP1. Although the induction of the TRAP1 complex was consistent with the pharmacological inhibition of ATP synthase across multiple cell lines, the variation in its protomer levels was not. While the TRAP1 complex is induced by inhibition of ATP synthase, it is reduced by inhibition of complexes I or III. Surprisingly, we found that inhibition of ATP synthase overrides the latter effect. This pharmacological epistasis experiment argues that ATP synthase is a primary TRAP1 interactor in the ETC. The opposite “perturbation” of OXPHOS, that is its stimulation by an inhibitor of lactate dehydrogenase, similarly promotes the formation of the TRAP1 tetramer. Thus, for reasons that remain to be elucidated, the “functional unit” of TRAP1 is sensitive to both an induction or a decline in OXPHOS.

*In toto*, although the precise molecular mechanism for how TRAP1 regulates OXPHOS remains to be uncovered, we know now that the overall levels of TRAP1 may not be correlated or relevant to OXPHOS regulation as previously thought [6]. It is really its tetrameric form that needs to be quantitated and structurally and functionally dissected in more detail to understand how TRAP1 contributes to regulating OXPHOS and mitochondrial homeostasis.

## Materials and methods

### Plasmids

The pcDNA3.1 (+) MTS-HA construct to direct all proteins to the mitochondrial matrix was generated by cloning the human TRAP1 mitochondrial targeting sequence between the EcoR1 site on the pcDNA3.1 (+) vector. All pcDNA3.1 (+) TRAP1-HA constructs including the truncation mutants were generated by cloning the human TRAP1 coding sequence into the pcDNA3.1 (+) MTS-HA construct. The TRAP1 coding sequence (without the MTS) was cloned into the XhoI restriction site after the TRAP1-MTS but before the HA-tag. The E115A/R402A and the ΔStrap mutants were subcloned from pPROEX HTb vectors into the XhoI site of the MTS-HA vector using the primers listed in Additional file 16: Table S9. The mitoEGFP construct was generated by cloning the EGFP coding sequence into the Xho1 site on the pcDNA3.1 (+) MTS-HA vector, exactly like TRAP1. mitoEGFP and all TRAP1 constructs with the pcDNA3.1 (+) MTS-HA vector have a C-terminal HA-tag. The TRAP1-FLAG and D158N-FLAG constructs were generated by cloning the TRAP1 coding sequence along with the C-terminal FLAG-tag between Kpn1 and Xho1 sites on the pcDNA3.1 (+) vector. For generating the TRAP1-GST construct, the TRAP1 coding sequence as a NheI-SalI fragment was joined to a SalI-EcoRI fragment carrying the GST coding sequence by insertion into the NheI-EcoRI sites of expression plasmid pcDNA3.1(+). The bacterial expression vector for the TRAP1 mutant E115A/R402A was generated from pTRAP1 [14] by site-directed mutagenesis using QuikChange (Agilent Technology). Sequences for all oligos are provided in Additonal file 16: Table S9. Note that for all TRAP1 point mutant, the numbering starts with the methionine of the MTS. The pHAGE-fEF1a-IZsGreen constructs used to stably express WT and E115A TRAP1 in MAFs were generated by cloning the respective sequences between the BamHI and NotI sites in plasmid pHAGE-fEF1a-IZsGreen (Additonal file 16: Table S9).

### Cell culture

HEK293T, HCT116, A549, UMUC3, MCF-7 and PC3 cells were obtained from American Type Culture Collection (ATCC, see Additional file 16: Table S9). MDA-MB-134 cells were obtained from Wilbert Zwart at the Netherlands Cancer Institute, Amsterdam. Unless specified otherwise, all cells were cultured at 37°C with 5% CO_2_in a standard incubator with Dulbecco’s Modified Eagle’s Medium (DMEM) glutamax, 4.5 g/l Glc and 1 mM Pyr (Thermo Scientific) supplemented with 10% fetal bovine serum (FBS), 100 u/ml penicillin and 100 µg/ml streptomycin.

### TRAP1 CRISPR/Cas9 knock outs

TRAP1 KO HEK293T and HCT116 cells were generated using CRISPR/Cas9 genome editing [61] as illustrated in Additional file 1: Figure S1A. The gRNA was designed using the online design tool by ATUM (https://www.atum.bio/eCommerce/cas9/input). The sense and antisense oligonucleotides for the selected gRNA construct (see Additional file 16: Table S9) were purchased (Microsynth), annealed and then inserted into the CRISPR/Cas9 vector PX459 (Addgene plasmid #48139) as previously described [61]. HEK293T and HCT116 cells were transiently transfected using polyethylenimine MAX (PEI) at a ratio of 1:3 of DNA to PEI. 48 hrs post transfection, the transfected cells were selected using 3-5 µg/ml puromycin until control non-transfected cells completely died. The remaining cells from the transfected population were allowed to grow in absence of puromycin until they formed visible foci. The cellular foci were then individually picked, subcultured and finally analyzed by immunoblotting with antibodies against TRAP1 to identify clones that were devoid of the protein. Three to five different KO clones for each cell line were frozen in liquid nitrogen. The A549 and UMUC3 TRAP1 KO clones were made using the all in one vector harboring a mCherry reporter (Genecopoeia, HCP200164-CG08-3; see Additional file 16: Table S9). The transfection procedure was similar to the one described for HEK293T and HCT116 cells, but the clonal isolation was performed with the mCherry reporter using FACS sorting under aseptic conditions. The sorted clones were subcultured and finally immunoblotted for TRAP1 to identify clones that were devoid of the protein.

### Cell culture for OCR experiments

Before any single carbon source OCR experiment, the cells were grown overnight in medium with the carbon source to be tested in order to acclimatize and to stabilize them metabolically. The carbon sources were added to DMEM lacking Glc, Pyr and Gln (A14430-01; see Additional file 16: Table S9) with 10% FBS, 100 u/ml penicillin and 100 µg/ml streptomycin as follows: (i) Glc only: 4.5 g/l glucose; (ii) Gln only: 2 mM glutamine; (iii) Pyr only: 1 mM sodium pyruvate; (iv) Gal and Pyr: 10 mM galactose, 1 mM sodium pyruvate.

### Flux assays

The mitochondrial OCR and ECAR were monitored *in vivo* in real-time using a Seahorse XF analyzer (XF^e^24, Agilent). Depending on the experiment, 6 x 10^4^ HEK293T or HCT116 cells were cultured overnight in custom XF24 microplates (poly-L-lysine coated) with either DMEM glutamax or DMEM (A14430-01) supplemented with the respective carbon sources. The standard assay medium used for all extracellular flux analyses and mitochondrial stress tests was unbuffered DMEM (SIGMA, D5030) without glucose, L-glutamine, sodium pyruvate, sodium bicarbonate, phenol red and FBS. Depending on the experiment, the D5030 medium was supplemented with the desired carbon source as indicated above. Prior to measurements, the cells were washed with and then incubated in unbuffered media (D5030) containing the respective carbon source in the absence of CO_2_ for 1 hr to acclimatize them to the assay medium. Following preincubation, basal OCR or ECAR were determined before recording mitochondrial stress test profiles by sequential injection of oligomycin, carbonyl cyanide-p-trifluoromethoxyphenylhydrazone (FCCP) and rotenone with antimycin in combination. For LDHi experiments, the LDHi (developed by the National Cancer Institute Experimental Therapeutics (NExT) Program) [62, 63] was injected first followed by an injection of oligomycin, rotenone and antimycin in combination to completely inhibit mitochondrial respiration.

For all assays involving transfected constructs, 2 x 10^5^ cells were first seeded in 6 well plates and allowed to grow overnight in DMEM glutamax. They were transfected on day 2 with 3 µg DNA using PEI for 6 hrs and further incubated overnight in DMEM glutamax. On day 3, 6 x 10^4^ transfected cells were seeded in polylysine-coated XF24 microplates and incubated in DMEM glutamax overnight. Real-time OCR and ECAR analyses were done as described above. For Gln only OCR analysis involving transfected constructs, the 6 x 10^4^ cells finally seeded for analysis on day 3 were incubated overnight in DMEM (A14430-01) supplemented with Gln.

### Glucose uptake and flow cytometry

The Glc uptake assay was performed with WT and KO HEK293T cells. On day one, 5 x 10^5^ cells were seeded and allowed to grow overnight in DMEM glutamax. On day 2, the cells were washed and incubated in DMEM (A14430-01) without any carbon sources for 1 hr to starve the cells of glucose before being incubated in DMEM supplemented with 150 µg/ml 2-NBDG. Cells were allowed to grow in this medium for 6 hrs. Cells were harvested by trypsinization, thoroughly washed in phosphate-buffered saline (PBS) and resuspended in 500 µl of PBS. Cells were initially analyzed using a BD FACsCaliber and its software CellQuest Pro. The final data analysis was done using the software FlowJo.

### Total metabolite and flux analysis using ^13^C-Gln

The metabolic flux analysis using ^13^C-Gln was performed by Human Metabolome Technologies, Inc. (https://humanmetabolome.com/en/targeted.html). Two biological replicates each of HEK293T and A549 cells were used for this experiment and grown in medium containing unaleblled Glc and Pyr, and ^13^C-labelled Gln (^13^C-Gln). Samples were prepared according to guidelines of the service provider from 5 x 10^6^ cells/ replicate and resuspended in 50 µl ultrapure water before measurements. The samples were analyzed using capillary electrophoresis time-of-flight mass spectrometry (CE-TOFMS, Agilent Technologies) in 2 modes to detect both anionic and cationic metabolites [64-66]. Detected peaks were then extracted using MasterHands ver. 2.17.1.11 to obtain m/z, migration time (MT) and peak area. Putative metabolites were assigned based on HMT’s target library and their isotopic ions on the basis of m/z and MT. Absolute quantitations were performed for the total amount of each detected metabolite.

### ATPase activity assay with the TRAP1 mutant E115A/R402A

#### Protein expression and purification

WT and TRAP1 mutant E115A/R402A were overexpressed in *Escherichia coli* BL21 (DE3)-RIL cells at 25°C following induction with 0.4 mM isopropyl β-D-1-thiogalactopyranoside at O.D._600_ ∼0.7. Cells were resuspended in buffer A (40 mM Tris-HCl pH 7.5, 400 mM KCl and 6 mM β-mercaptoethanol) and lysed using a microfluidizer. The cleared lysate was loaded onto a pre-equilibrated Ni-NTA agarose column (Qiagen) and washed with buffer A supplemented with 30 mM imidazole. Bound protein was eluted using a linear gradient from 30 to 500 mM imidazole in buffer A. Peak fractions were pooled, mixed with His_6_-TEV protease, and dialyzed against buffer B (25 mM Tris-HCl pH 8.0, 100 mM NaCl and 6 mM β-mercaptoethanol). The liberated His-tag and His-TEV were removed by reapplying the sample to a Ni-NTA agarose column. Ammonium sulfate to a final concentration of 0.5 M was added to the flow-through, which was loaded onto a pre-equilibrated TOYOPEARL Butyl 600M column (Tosoh Bioscience), eluted using a linear gradient of 0.5 to 0 M ammonium sulfate in buffer C (25 mM Tris-HCl pH 8.0 and 6 mM β-mercaptoethanol), and dialyzed against buffer D (25 mM Tris-HCl pH 7.5, 100 mM KCl, and 6 mM β-mercaptoethanol).

#### ATPase assay

ATPase activities were determined with recombinant protein at 10 µM at 30°C in 30 mM HEPES/KOH pH 7.5, 50 mM KCl, 5 mM MgCl_2_, 2 mM DTT and 2 mM ATP by measuring the amount of inorganic phosphate released after 30 min using the malachite green calorimetric assay [67].

### Isolation of mitochondria

Mitochondria were isolated from cells grown in large 15 cm dishes to approximately 95% (not 100%) confluency using a protocol adapted from Martinou and coworkers [68]. Briefly, cells were trypsinized, washed and pelleted in ice-cold PBS (1,000 rpm, 5 min), and then re-suspended in 2 ml ice-cold MB buffer (10 mM Hepes pH 7.5, 210 mM mannitol, 70 mM sucrose, 1 mM EDTA) and manually homogenized using a Dounce homogenizer (50 times per sample). The homogenate was centrifuged at 2,000 xg for 10 min to pellet nuclei and cell debris. The supernatant was spun again at 16,000 xg for 10 min. The resulting brown pellet contained mitochondria and was rigorously washed six times with ice-cold MB buffer by resuspending and centrifugation at 16,000 g for 10 min.

### TRAP1 IPs

For all IP experiments, the mitochondria isolated from cells expressing various TRAP1 constructs were resuspended in lysis buffer (10 mM Tris-HCl pH 7.5, 50 mM NaCl, 1mM EDTA, 1mM DTT, 10% glycerol, 10 mM sodium molybdate, 0.1% Triton X-100 and protease inhibitor cocktail (A32965, Thermo Scientific)) and lysed by sonication (35 cycles of 30 sec) using a Bioruptor (Diagenode). For all IPs, 1 mg clarified mitochondrial lysate was incubated overnight with 3 µg anti-HA antibody at 4°C on a spinning rotor. The following day, 50 µl of Dynabeads-Protein G (10009D, Thermo Scientific) were added to the antibody-lysate mix and incubated at 4°C on a spinning rotor for 3 hrs. Following incubation, the Dynabeads were washed four times with lysis buffer. The proteins were eluted with NuPAGE sample buffer supplemented with 10 mM DTT.

### TRAP1 mutant IP-MS analysis and comparison

The TRAP1 mutant IP-MS analysis was performed by Poochon Scientific (https://www.poochonscientific.com/services/protein-identification/) with three biological replicates per sample and two replicates for controls. Briefly, 2 x 10^6^ HEK293T cells were seeded in 15 cm dishes, grown and transfected with various constructs using the Jetprime transfection reagent at 70% confluency. 24hrs after transfection, mitochondrial lysate preparation and IPs were performed as described above. 30 µl of the total IP sample for each IP (two controls and triplicates for the mutants) were run on a 4-12% gradient SDS-PAGE followed by in-gel trypsin digestion and LC/MS/MS analysis. The LC/MS/MS analyses of samples were carried out using a Thermo Scientific Q-Exactive hybrid quadrupole-orbitrap mass spectrometer and a Thermo Dionex UltiMate 3000 RSLCnano system. For each LC/MS/MS run, the tryptic peptide mixture was loaded onto a peptide trap cartridge set to a flow rate of 5μl/min. The trapped peptides were eluted onto a reversed-phase PicoFrit column (New Objective, Woburn, MA) using a linear gradient of acetonitrile (3-36%) in 0.1% formic acid. Eluted peptides from the PicoFrit column were then ionized and sprayed into the mass spectrometer, using a Nanospray Flex Ion Source ES071 (Thermo Scientific). For protein identification, two raw MS files from two LC/MS/MS runs for each sample were analyzed using the Thermo Proteome Discoverer 1.4.1 platform (Thermo Scientific, Bremen, Germany) for peptide identification and protein assembly. Database searches against the public human protein database obtained from the NCBI website were performed based on the SEQUEST and percolator algorithms through the Proteome Discoverer 1.4.1 platform. The minimum peptide length was specified to be five amino acids. The precursor mass tolerance was set to 15 ppm and the fragment mass tolerance was set to 0.05 Da. The maximum false peptide discovery rate was specified as 0.01. Finally, the estimation of relative protein abundance was based on PSMs. For further comparison of relative abundance of interacting proteins for a particular mutant or for WT TRAP1, all data were normalized to 100 PSMs for the immunoprecipitated TRAP1 protein in a given replicate.

### Stable isotope labeling by amino acids in cell culture (SILAC)

SILAC was performed as follows. As culture medium, DMEM deprived of lysine and arginine was used together with dialyzed fetal bovine serum (10 kDa cutoff). For light medium, L-lysine-2HCl was added to a final concentration of 146.2 mg/l and L-arginine-HCl was added to a final concentration of 84 mg/l. For heavy medium, L-lysine-2HCl (^13^C_6_, ^15^N_2_) was added to a final concentration of 181.2 mg/l and L-arginine-HCl (^13^C_6_, ^15^N_4_) was added to a final concentration of 87.8 mg/l. Heavy and light SILAC labeling was achieved by culturing UMUC3 cells in the respective media for 5 cell doublings (replenishing media every 2-3 days). Care was taken to maintain the UMUC3 cell cultures in their log phase of growth. Separate stable cultures of WT and TRAP1 KO UMUC3 cells were established in both heavy and light DMEM. After 5 cell doublings, heavy labeling efficiency was determined to be >95%. At this point, comparative steady-state protein expression in both heavy-labeled KO cells and light-labeled WT cells (or vice versa) was performed in triplicate samples (biological replicates) by the Mass Spectrometry Section of the Collaborative Protein Technology Resource (Center for Cancer Research, National Cancer Institute, Bethesda, MD).

### LFQ MS analysis

Three biological replicates of 9 x 10^6^ WT and KO HEK293T and HCT116 cells grown in different carbon source cocktails (Glc + Pyr + Gln, Gal+ Pyr and Gln only) were pooled together and lysed in FASP lysis buffer (100 mM Tris-HCl pH 7.5, 4% SDS, 10 mM TCEP) at 95 °C for 5 min followed by centrifugation at 14,000 g for 10 min.100 µg of each clarified sample were digested by the FASP method [69]. 50 µg of the resulting peptide mixtures were desalted on Waters SEP-PAK C18 micro elution plates and eluted with 100 μl of 40% acetonitrile, 0.1% formic acid. 6 μl of the eluate were used for the MS analysis using a Thermo Scientific Q-Exactive hybrid quadrupole orbitrap fusion mass spectrometer. Data analysis was done using MaxQuant and Perseus.

### Native-PAGE

30 µg total mitochondrial protein extracts were resolved on 6% or 8% Tris-glycine clear native gels. The pH values for the stacking and separating parts of the gels, and for the running buffer were 8.8 and 6.8, and 8.3, respectively. Gels were run at 80 V for 5-6 hrs at 4°C. The resolved proteins were transferred onto nitrocellulose membranes overnight at 30 V at 4°C. TRAP1 complexes were revealed by immunoblotting with an anti-TRAP1 antibody (BD Biosciences).

### Drug treatments

2 x 10^6^ HEK293T, HCT116, MCF-7, MDA-MB-134 or PC3 cells were seeded and grown to 90-95% confluency in 15 cm plates. Depending on the experiment, the cells were treated with 10 µM oligomycin (complex V inhibitor), rotenone (complex I inhibitor) or antimycin (complex III inhibitor) for 2, 4, 6 or 8 hrs in medium containing Glc, Pyr and Gln as carbon sources. Following drug treatments, mitochondrial extracts were prepared and native PAGE run as described above. For LDH inhibition, 5 µM of the LDHi was used for 2, 4 and 6 hrs.

### TRAP1-GST pulldown

2 x 10^6^ HEK293T cells were seeded in 15 cm dishes, grown and transfected with expression vectors for TRAP-GST and GST using the Jetprime transfection reagent at 70% confluency. 24 hrs after transfection, mitochondrial lysates were prepared in lysis buffer (10 mM Tris-HCl pH 7.5, 50 mM NaCl, 1mM EDTA, 0.1% Triton X-100, 1mM DTT, 10% glycerol, 10 mM sodium molybdate, protease inhibitor cocktail (A32965, Thermo Scientific)) as described before. 1 mg clarified mitochondrial lysates prepared in lysis buffer was incubated overnight with 50 µl glutathione-conjugated magnetic agarose beads (Thermo Scientific) at 4°C on a spinning rotor. The beads were washed four times with the same buffer and the proteins were eluted at room temperature in the same buffer supplemented with 80 mM reduced glutathione. The eluted samples were immediately run on a 6% clear native gel and processed for MS as illustrated in Additional file 12: Figure S5a.

### MS analysis of oligomeric TRAP1 complex

The TRAP1 complexes from the GST pulldowns were visualized on the native gels by staining with coommassie brilliant blue (CBB G-250) followed by sequential destaining. The portion of the gel containing the stained TRAP1-GST complex was extracted as shown in Additional file 12: Figure S5a (equivalent position on the gel was extracted for controls; see Additional file 12: Figure S5a). The extracted gel slices were first reduced with DTT and then alkylated with iodoacetamide. Next, the samples were trypsin digested. The digested peptide mixture was then concentrated and desalted using C18 Zip-Tip. The desalted peptides were reconstituted in 20 μl of 0.1% formic acid. From this, 18 μl of peptides were analyzed by LC/MS/MS using a Thermo Scientific Q-Exactive hybrid quadrupole-orbitrap mass spectrometer and a Thermo Dionex UltiMate 3000 RSLCnano System as described above for TRAP1 IP-MS. Proteins in the oligomeric TRAP1 complex were determined by filtering the data for proteins with a high number of unique peptides and cross-referencing with the GST control to eliminate overlapping proteins as illustrated in Additional file 12: Figure S5b.

### Q-PCR analysis

2 x 10^5^ WT HEK293T cells were seeded in 6 well plates overnight. On day 2, one set was transfected with a HIF1α expression vector [70] (see Additional file 16: Table S9) using the Jetprime transfection reagent. On the same day, one set was exposed to hypoxia (1% O_2_, overnight) and the third set was left in normoxia. On day 3, each set was collected and analyzed by quantitative reverse-transcription PCR (RT-PCR) with specific primers (Additional file 16: Table S9). Briefly, RNA was isolated with the acid guanidinium thiocyanate-phenol-chloroform method [71]. 500 ng RNA was used for reverse transcription using random primers and the GoScript master mix according to the manufacturer’s instructions (Promega). Quantitative real-time PCR was used to examine the expression levels of *TRAP1* and *HIF1A* with *GAPDH* as the reference gene.

### Statistical analyses

Data analysis was primarily performed using Graphpad Prism 8, Perseus (MS) and Microsoft Excel. The differences between various groups was analyzed with a two tailed Students t-test. Until specified, the error bars represent the standard error of the mean with *p<0.05, **p<0.01, and ***p<0.001 denoting the difference between the means of two compared groups considered to be statistically significant. Each real-time OCR tracing profile shown represents a cumulative plot of three technical replicates per cell type.

## Additional files

**Additional file 1: Figure S1.**
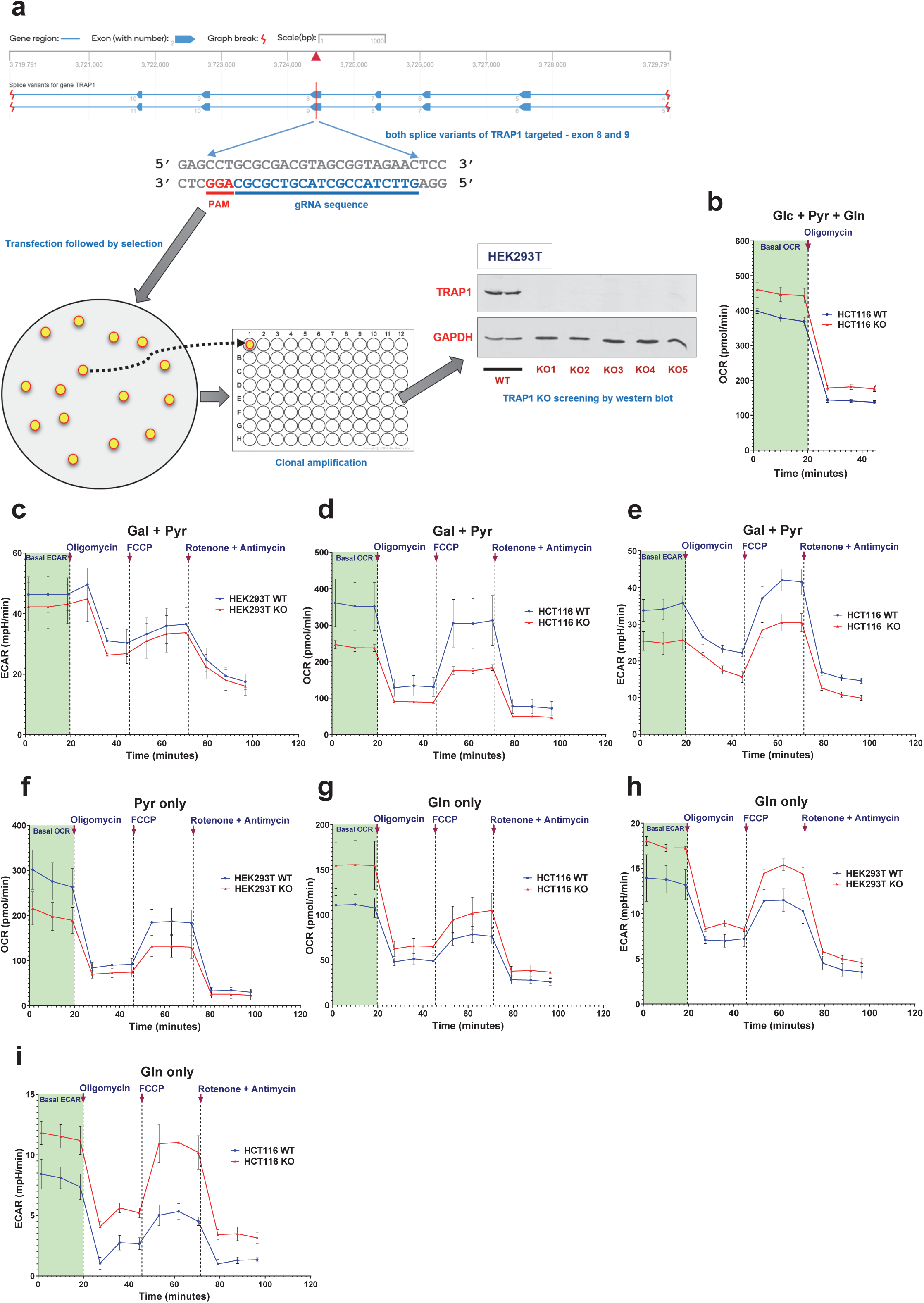
Generation of TRAP1 KO cells and additional metabolic profiling. (a) Workflow for the generation of CRISPR/Cas9-mediated TRAP1 KO clones. Unlike HEK293T and HCT116 clones, A549 and UMUC3 TRAP1 KO clones were isolated by fluorescence-activated cell sorting using a vector allowing mCherry expression (see Additional file 16: Table S9). (b) OCR traces of WT and KO HCT116 cells with Glc + Pyr + Gln as carbon sources. (c-i) OCR and ECAR traces of WT and KO HEK293T or HCT116 cells with different primary carbon sources.

**Additional file 2: Figure S2.**
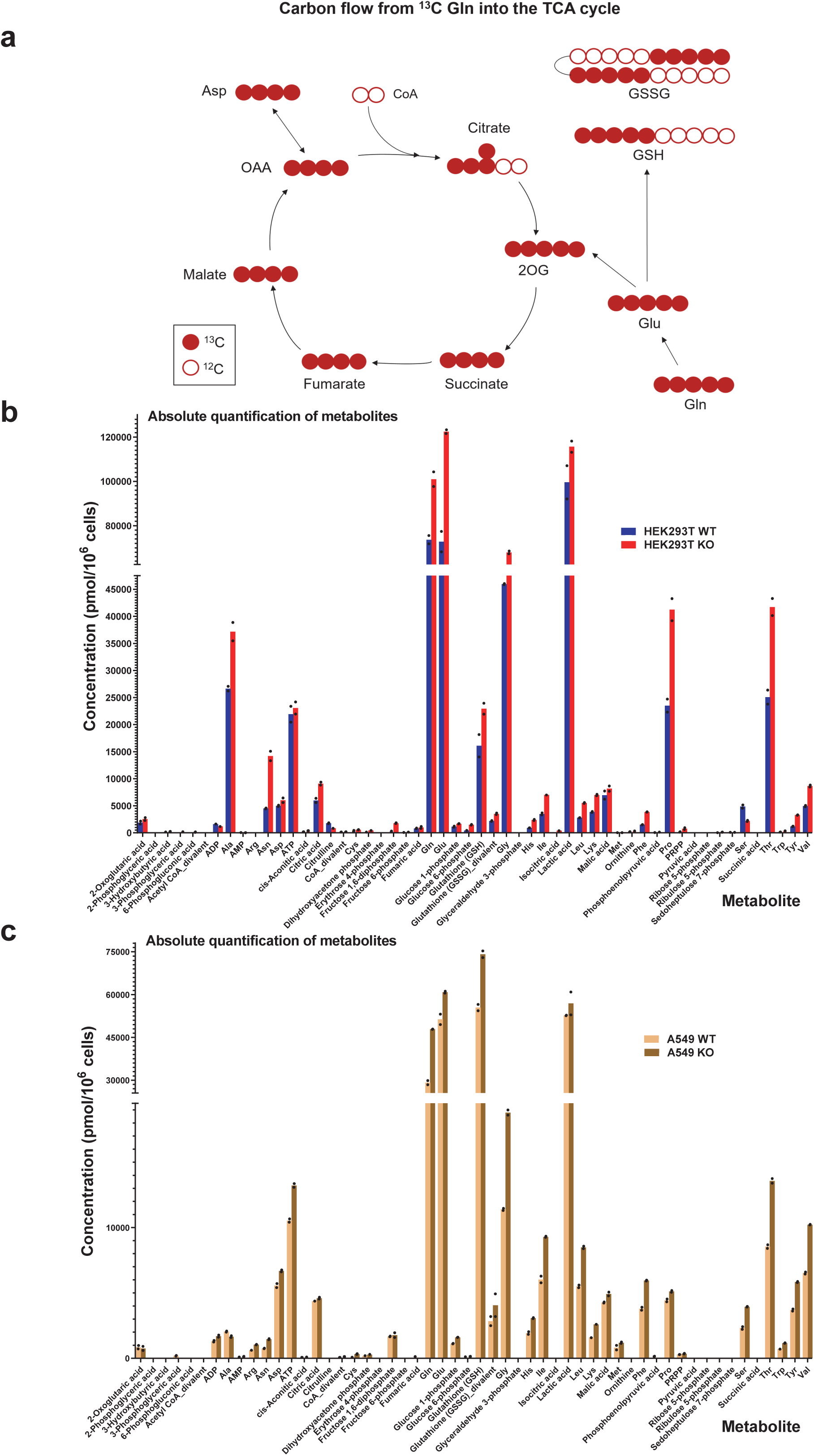
Carbon flux and total quantitation of target metabolites. (a) Schematic metabolic map showing the flow and distribution of ^13^C atoms in metabolites of the TCA cycle when cells consume ^13^C-Gln. Note that most of these metabolites traced with ^13^C-Gln were found to be upregulated in TRAP1 KO cells. (b, c) Total quantitation of target metabolites in WT and KO HEK293T and A549 cells. Note that this is “total” quantitation and should not be confused with ^13^C tracing. Total quantitation must be combined with the information provided in Additional file 4: Table S2 to infer metabolites with increased ^13^C incorporation. Data points on bar graphs indicate metabolite concentration per 10^6^ cells from each biological replicate (n = 2).

**Additional file 3: Table S1.** Quantitative estimation of target metabolites in HEK293T and A549 cells.

**Additional file 4: Table S2.** Quantitative ^13^C tracing in target metabolites in HEK293T and A549 cells.

**Additional file 5: Figure S3.**
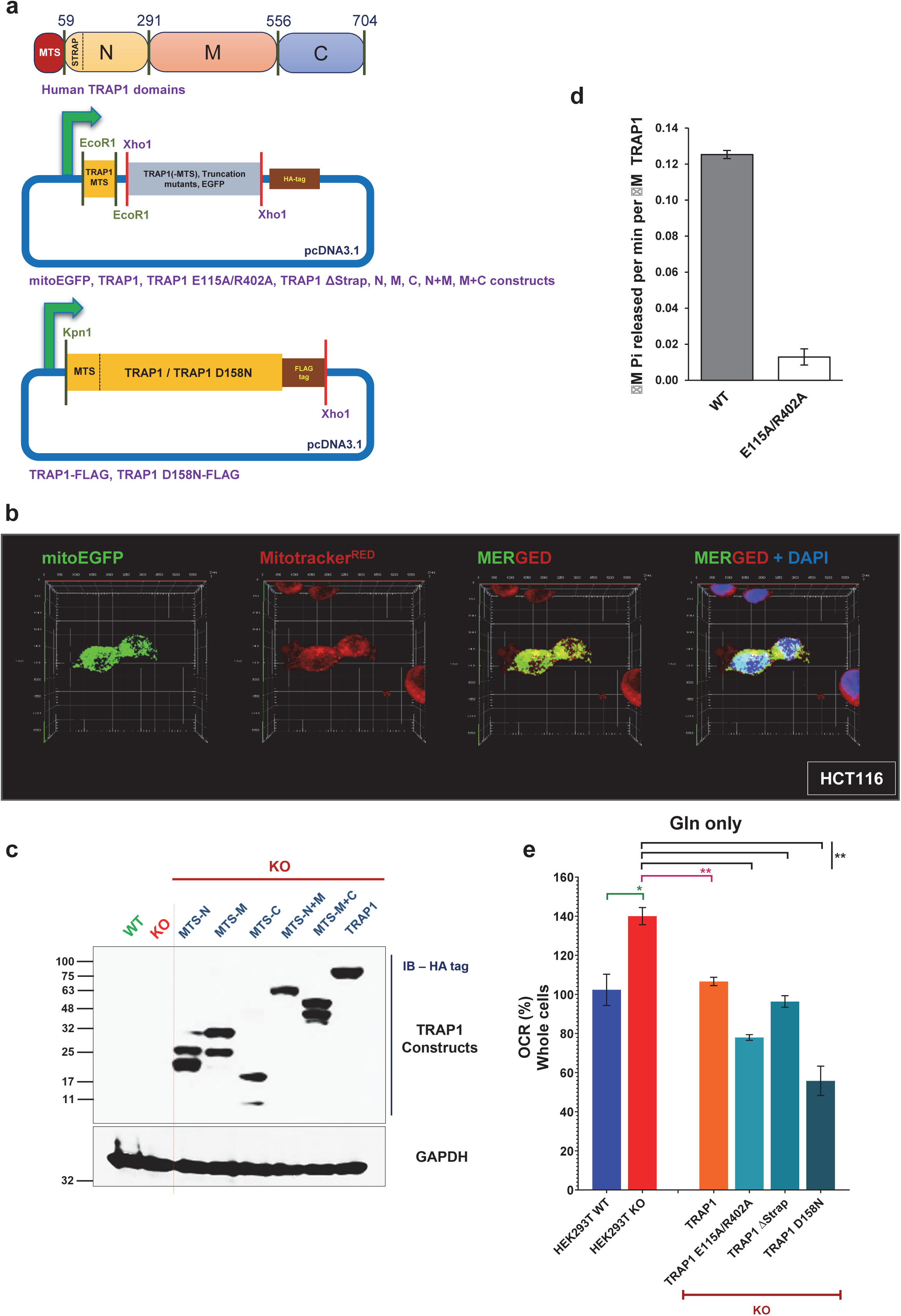
TRAP1 truncation and point mutants. (a) Schematic representation of the constructs for expression of mitochondrially targeted TRAP1 and EGFP. (b) Fluorescence micrographs showing proper targeting of mitoEGFP to mitochondria. Mitochondria are revealed with Mitotracker^RED^. (c) Expression analysis of TRAP1 truncation mutants by immunoblotting with an antibody to their HA-tag. (d) ATPase activity assay for the TRAP1 double mutant E115A/R402A. (e) Quantitation of basal respiration rates in WT versus KO HEK293T cells expressing the indicated proteins. Note that all ATPase mutants can rescue the KO phenotype to WT levels.

**Additional file 6: Figure S4.**
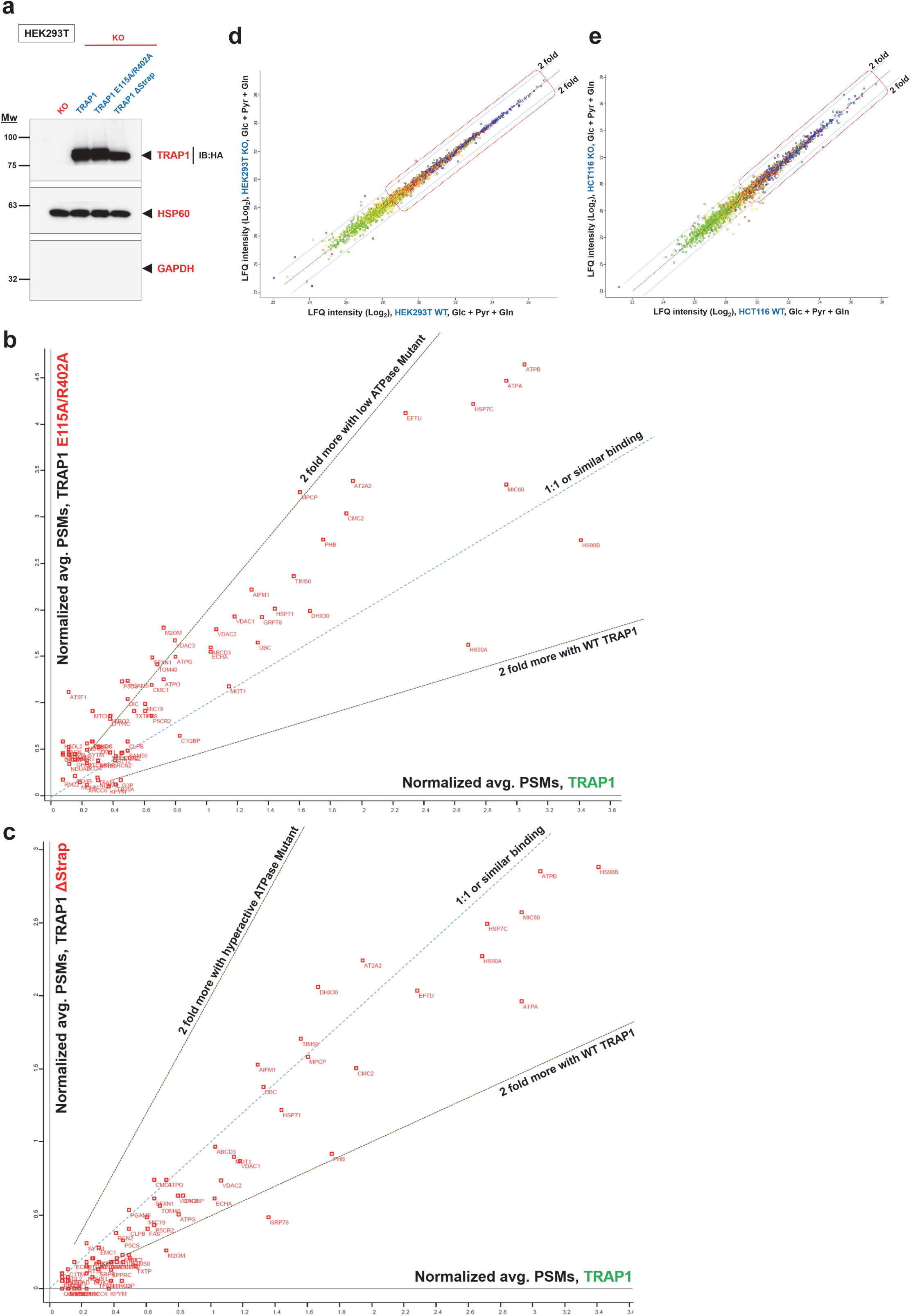
Analysis of the whole cell proteome and TRAP1-associated proteins. (a) Control immunoblot performed to check TRAP1 WT and mutant expression in the KO cells used for the IP-MS experiments. (b, c) Comparative relative abundance of proteins immunoprecipitated with the indicated TRAP1 ATPase muatnts or WT TRAP1. The scatterplot was generated as mentioned in the legend to Fig. 4a. (d, e) Scatter plots comparing the levels (LFQ intensities) of the 2660 high confidence proteins between WT and KO HEK293T or HCT116 cells.

**Additional file 7: Table S3.** List of all identified proteins pulled down with TRAP1 using an IP-MS analysis with WT TRAP1, and the TRAP1 mutants E115A/R402A and ΔStrap.

**Additional file 8: Table S4.** List of high confidence TRAP1 interacting proteins (from Additional file 10: Table S3) filtered for mitochondrial localization and a minimum of 4 or more identified unique peptides (with a few exceptions).

**Additional file 9: Table S5.** List of mitochondrial proteins identified in the SILAC analysis comparing WT to TRAP1 KO UMUC3 cells. Note that only those proteins were considered that were identified and quantitated in all three replicates.

**Additional file 10: Table S6.** Complete list of proteins identified in whole cell LFQ MS analysis to compare WT to TRAP1 KO HEK293T and HCT116 cells.

**Additional file 11: Table S7.** List of high confidence proteins identified in whole cell LFQ analysis to compare WT to TRAP1 KO HEK293T and HCT116 cells. The 4578 proteins from Additional file 10: Table S6 were reduced to 2660 by selecting only those with at least 7 identified unique peptides in the LFQ analysis.

**Additional file 12: Figure S5.**
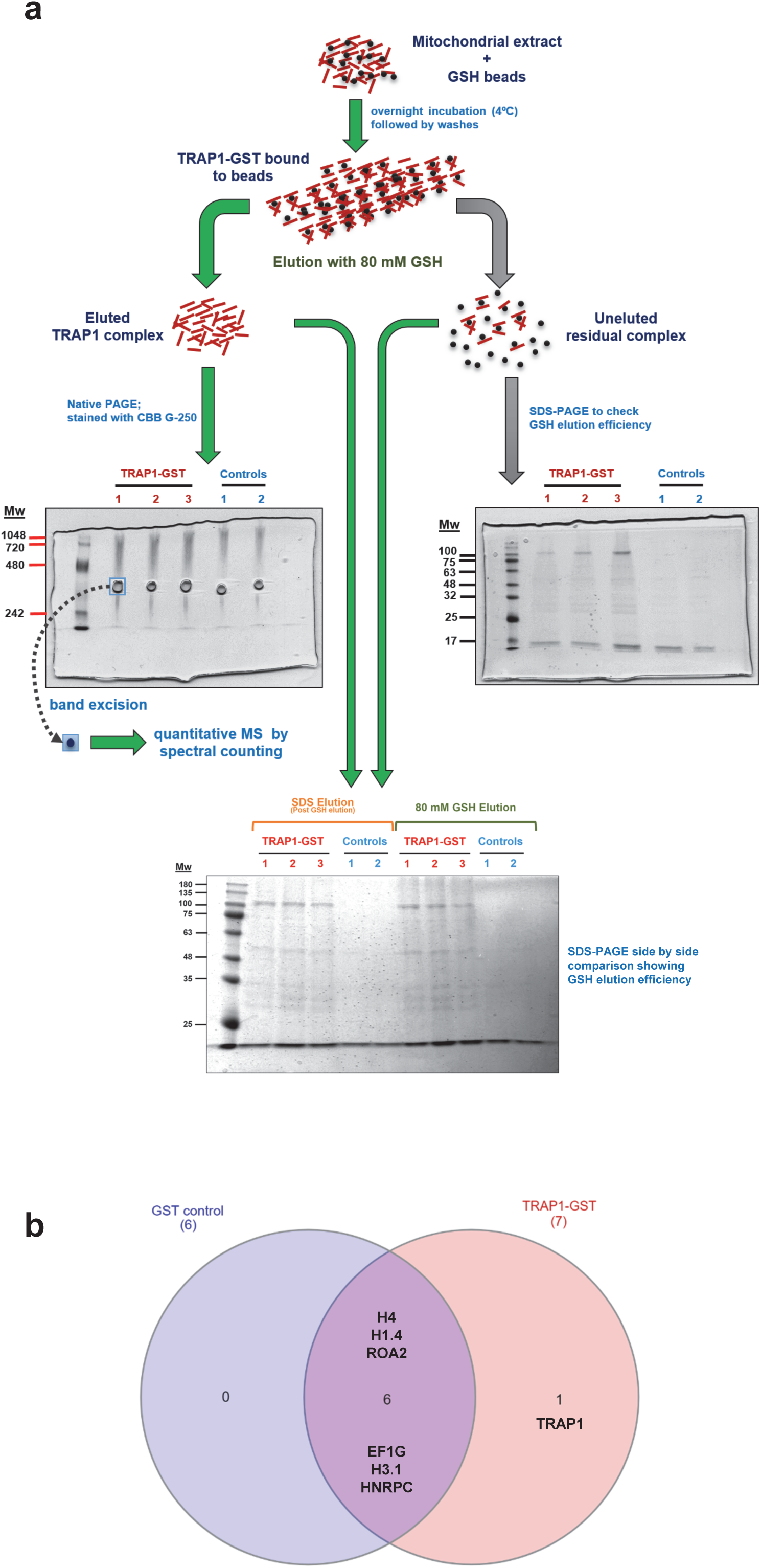
An extension of Figure 5 showing TRAP1-GST pulldown MS strategy and analysis. (a) TRAP1-GST pulldown strategy. (b) Venn diagram of the proteins identified by the MS analysis. Note that TRAP1 peptides are the only unique ones in the TRAP1-GST pulldown samples compared to the GST controls.

**Additional file 13: Table S8.** TRAP1 complex MS analysis.

**Additional file 14: Figure S6.**
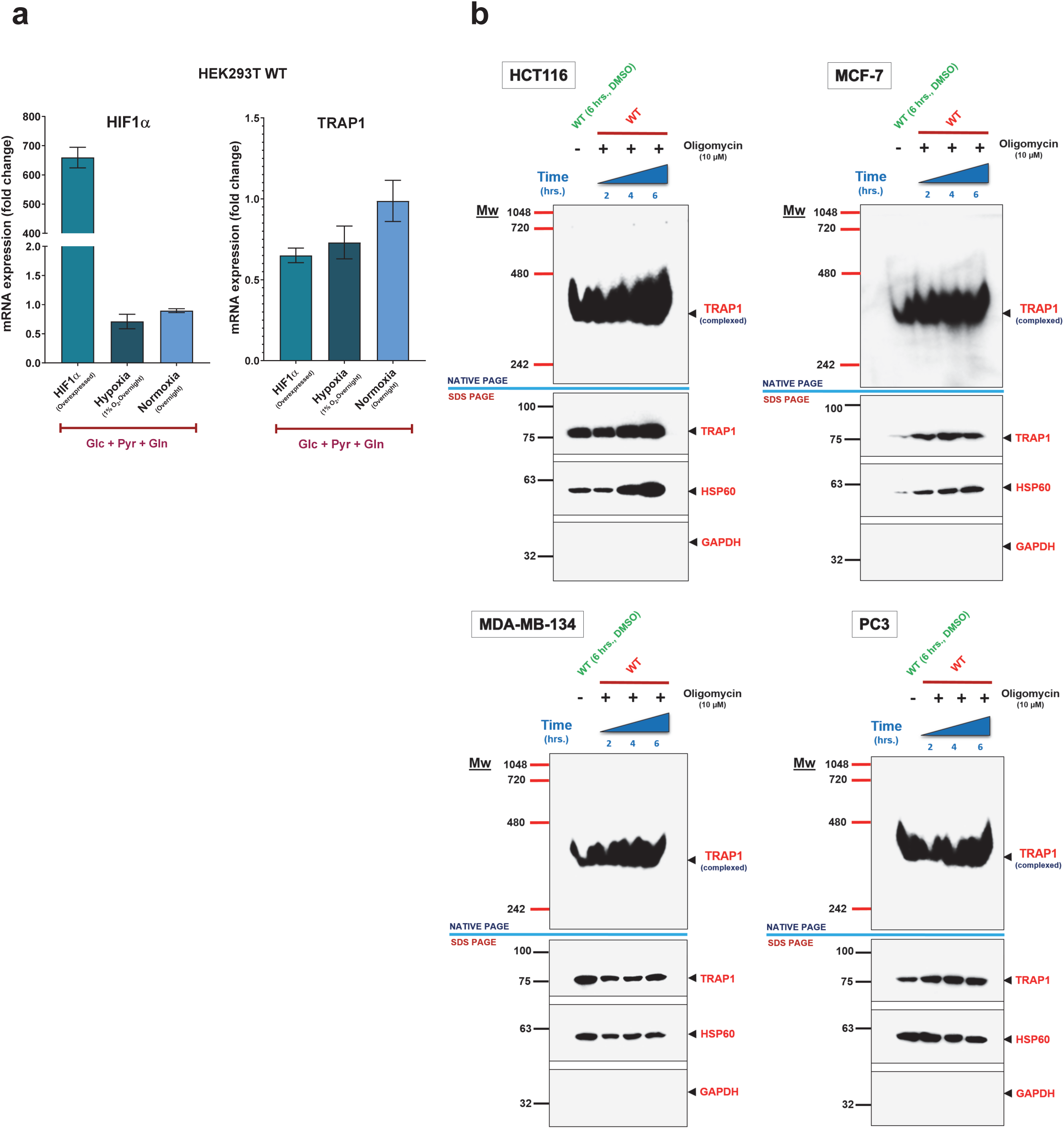
TRAP1 is not induced by HIF1α and the TRAP1 complex is ubiquitous. (a) Quantitative RT-PCR analysis of the mRNA levels for HIF1α and TRAP1. All data are reported as means ± SEM (n = 3). (b) Analysis of TRAP1 complexes from indicated cell lines by native PAGE and SDS-PAGE.

**Additional file 15: Figure S7.**
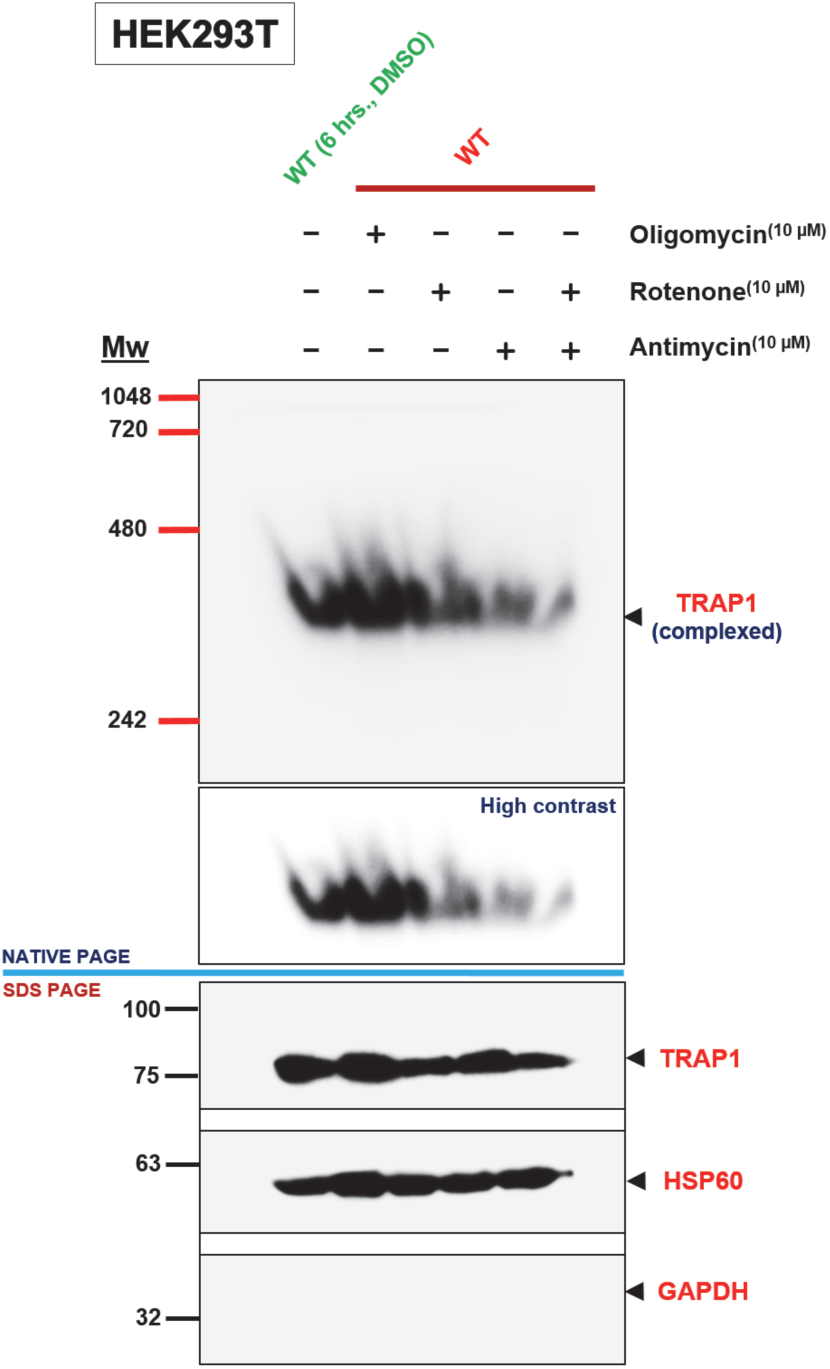
Differential effects of drug treatments on the TRAP1 complex.

**Additional file 16: Table S9.** List of reagents and resources.

## Acknowledgements

We thank Jean-Claude Martinou for invaluable discussions regarding this project.

## Availability of data and materials

All data generated during this study are included in either the published article or its Additional files.

## Authors’ contributions

A.J. conceived the study, designed and performed experiments, analyzed the data, prepared figures, and wrote the manuscript. J.D. designed and performed the quantitative metabolic flux and SILAC experiments. N.G. performed and analyzed the Q-PCR data. J.L. and F.T.F.T. purified the TRAP1 E115A/R402A mutant and performed and analyzed its ATPase activity. G.S. helped in the analysis of SILAC data. K.B. helped with the analysis of proteomics data. L.N. contributed to understanding metabolic dynamics, designing and supervising experiments, and to writing the manuscript. D.P. supervised the work, contributed to designing experiments, and wrote and critically edited the manuscript. All authors contributed to the overall editing of the manuscript.

